# Asymmetric Boost Responses to an HIV-1 Vaccine: Boost-site Controls Secondary B-cell Fates

**DOI:** 10.64898/2026.08.11.744292

**Authors:** John S. Barber, Keisuke Tonouchi, Chen-Hao Yeh, Madison Berry, Helene F. Kirshner, Kevin Wiehe, Amanda Eaton, David C. Montefiori, Ming Tian, Frederick W. Alt, Kevin O. Saunders, George M. Shaw, Barton F. Haynes, Garnett Kelsoe

**Affiliations:** Division of Allergy and Immunology, Department of Pediatrics, Duke School of Medicine, Durham, NC 27710, USA; Department of Integrative Immunobiology, Duke School of Medicine, Durham, NC 27710, USA; Duke Human Vaccine Institute, Duke School of Medicine, Durham, NC 27710, USA; Department of Surgery, Duke School of Medicine, Durham, NC 27710, USA; Howard Hughes Medical Institute, Boston Children’s Hospital, Boston, MA 02115, USA; Program in Cellular and Molecular Medicine, Boston Children’s Hospital, Boston, MA 02115, USA; Department of Genetics, Harvard Medical School, Boston, MA 02115, USA; Department of Molecular Genetics and Microbiology, Duke School of Medicine, Durham, NC 27710, USA; Departments of Medicine and Microbiology, Perelman School of Medicine, University of Pennsylvania, Philadelphia, PA 19104, USA; Duke Department of Medicine, Duke School of Medicine, Durham, NC 27710, USA

## Abstract

Induction of broadly neutralizing antibody (bnAb) responses remains a central challenge to designing efficacious HIV vaccines. *Lineage design* strategies, in which bnAb precursors are guided via iterative immunizations to their mature forms, depend on high levels of somatic mutation and the recall of antigen-specific B cells. Recent studies have identified spatial context as an important determinant of boosting efficacy, but the application of this to HIV vaccines and the underlying mechanisms are incompletely understood. Here, using mice harboring a V3-glycan bnAb lineage precursor knock-in combined with lineage-tracing and single-cell analyses, we show that persistent germinal centers (GCs) support continued affinity maturation of founder clones and ipsilateral boosting preferentially engages these lineages in secondary GCs. In contrast, contralateral boosting predominantly recruits naïve B cells and memory B cells not directed towards the immunizing antigen. The few memory cells recruited at this site were biased towards a plasma cell fate. Finally, we identify disfavored mutational trajectories within the V3-glycan bnAb lineage, revealing intrinsic constraints on bnAb lineage evolution.

## Introduction

Effective HIV-1 vaccines must elicit potent and durable broadly neutralizing antibody (bnAb) responses. This goal remains elusive, in part a consequence of the improbable but critical features of bnAbs including: rare precursor B-cell frequencies^1; 2^; uncommonly high frequencies of somatic V(D)J mutation^3–7^; a necessity for critical but improbable mutations^8^, and a poly- or autoreactivity normally suppressed by immune tolerance mechanisms^9^. These unusual but characteristic features of bnAbs have resulted in vaccine development strategies to direct the somatic evolution of HIV-1 reactive B cells along the rare evolutionary trajectories leading to bnAb generation^10–12^.

These new vaccine strategies are informed by the recovery of specific bnAb lineages from infected patients and seek to replicate bnAb B-cell development by the design of sequential immunogens that select for critical mutations ensuring neutralization potency and breadth^10; 13^. In brief, this *lineage design* vaccine strategy is iterative with an initial immunization targeted for the activation and proliferative expansion of the rare, unmutated ancestors (UCAs) of bnAb lineages followed by serial boosters aimed at selecting for critical mutations. In this way, bnAb lineage BCR evolution may be guided towards neutralization breadth and potency. Optimism for the potential utility of *lineage design* vaccination has grown based on a number of studies showing the potential of designed immunogens to activate precursor BCRs in animal models^14–18^ and humans^19; 20^. Given the challenges in eliciting bnAbs, a critical prerequisite of *lineage design* strategies is that they promote somatic mutation across successive rounds of immunization within a single clonal lineage. While the existence of long-lived germinal centers in response to complex antigens has been known^21^, interest has grown recently regarding their potential in protective responses to vaccination and infection^22^. Several studies have concluded that persistent germinal centers (GCs) are mostly populated by recent immigrant B cells, often with low avidity for the eliciting immunogen, and that these cells may directly compete with memory B-cells for affinity maturation^23; 24^. This competition could limit the step-wise development of B-cell lineages with critical mutations. These studies, however, also identified a small B-cell population in persistent GCs characterized by a high degree of somatic mutation and strong antigen affinity; this population likely represents the progeny of successful GC founder B cells. Tracking the founder B-cell population separately from newly recruited immigrants enables proper assessment of the ability to acquire key mutations.

Another potential weakness in *lineage design* vaccine strategy is the requirement that each successive round of boost immunization acts efficiently to re-activate the memory B cell (Bmem) populations selected and expanded by prior vaccinations. This presents a critical challenge as Victora has shown that recruitment of Bmem into recall germinal centers (GCs) is very inefficient^25^. Instead, on antigen re-exposure, the great majority of Bmem appear to proliferate and differentiate into long-lived plasma cells^26–28^. This differentiation bias may be compounded further as several studies have concluded that high-affinity B cells are preferentially fated for plasmacytic differentiation^29; 30^. If so, serial immunizations could result in the specific depletion of the very high-affinity B cells that *lineage design* vaccination is designed to produce.

That the functional properties of local and systemic immune memory differ is now well recognized. Resident local Bmem are established after pulmonary influenza infection in mice and these cells differ from systemic Bmem in their capacity to differentiate into plasma cells^31^. Our recent work extended this property to secondary lymphoid organs by demonstrating the superiority of local (ipsilateral) boosts in supporting Bmem reentry into GCs compared to distal (contralateral) boosts with influenza hemagglutinin antigen^32^. Dhenni et al.^33^ further demonstrated that Bmem in draining lymph nodes, but not in non-draining lymph nodes, reside within the subcapsular sinus niche where sinus macrophages maintain them, and that Bmem from distal sites exhibit a bias toward plasma cell differentiation. Thus, although the prevailing paradigm suggests that Bmem fate is determined primarily by intrinsic properties^34^, there is increasing recognition of the importance of *where* these Bmem encounter antigen.

Here, we characterize in detail primary and secondary B-cell responses to an HIV stabilized envelope (Env) trimer (SOSIP) vaccine immunogen by the conditional marking of early, primary GC cells and their progeny. We aimed to determine whether persistent GCs support the development of bnAb lineage clones and whether the location of a homologous boost immunization is relevant for determining Bmem fate^31; 35^. To address these questions, we paired a S1pr2-driven GC fate-mapping system^36^ with mice harboring a BCR knock-in of the germline progenitor of the DH270 bnAb lineage. We show that, although GC B cell numbers decrease over time, a persistent population of marked, founder B cells remain and accumulate V(D)J mutations, including combinations of improbable mutations critical for bnAb activity^8; 13^. These marked, persistent B-cell lineages are efficiently expanded in secondary GC responses by ipsilateral, but not contralateral, boosts. In contrast, contralateral boosts reactivated marked progeny of early GC B cells but strongly skewed their proliferation and differentiation towards plasmacyte fate. Correlated with the asymmetric outcomes of ipsilateral and contralateral boosting was the retention of S1pr2-marked Tfh cells at sites of primary immunization and disproportionate representation of marked B cells that did not harbor the knock-in BCR at contralateral boost sites. These observations reveal a spatial component to secondary responses with important implications for vaccine design.

## Materials and Methods

### Mice and Immunizations

Female C57BL/6 mice were obtained from the Jackson Laboratory. DH270 UCA VH and VL KI mice were described previously (Saunders et al., 2019). S1pr2-ERT2Cre-tdTomato^36^ mice were provided by Dr. Tomohiro Kurosaki. F1 mice from DH270 UCA VH and VL KI and S1pr2-ERT2Cre-tdTomato breeders were used for all described experiments unless otherwise specified. All mice were maintained under specific pathogen-free conditions at the Duke University Animal Care Facility. Eight to twelve-week-old mice were immunized with 10 μg of 10.17DT-NP^15^ in the footpad of the right hind leg. Six days after immunization, tamoxifen (12.5) mg was administered per os. Eight to nineteen weeks later, cohorts of mice were boosted with 10 μg of 10.17DT-NP in the hock ipsilaterally (right hind leg) or contralaterally (left hind leg). All antigens were mixed with Alhydrogel® adjuvant 2% (InvivoGen, final concentration of 1%) before immunizations. LNs draining the site of the most recent immunizations (right popliteal LNs for no-boosts and ipsilateral boosts, and left popliteal LNs for contralateral boosts) were analyzed. Sera were collected at various timepoints throughout experiments as indicated. All experiments involving animals were approved by the Duke University Institutional Animal Care and Use Committee.

### Production of immunogen

CH848 10.17DT SOSIP gp140 trimers were produced as previously described ^15^. Briefly, chimeric SOSIP trimers were expressed in FreeStyle 293 cells with furin co-transfection, purified by PGT145 affinity chromatography, and further purified by size-exclusion chromatography.

### Flow Cytometry

Single-cell suspensions from recovered LNs were dispersed by gentle disruption between glass slides and suspended in 10% DMEM supplemented with 10% FCS, 55 µM 2-mercaptoethanol, 100 units/ml penicillin, 100 μg/ml streptomycin and additional 2 mM L-glutamine (all Invitrogen). Suspended cells were then pre-treated with the mixture of anti-CD16/CD32 Ab (2.4G2, BD) and rat IgG (Invitrogen) on ice to reduce unspecific interactions between cells and labeling Abs. After 30 minutes, cells were stained with fluorophore-conjugated Abs for an additional 30 minutes. Abs used in this study are as follows: Anti-GL7 FITC (BD Biosciences), anti-IgM PerCP eF710 (II/41, eBioscience), anti-CD38 PE-Cy7 (90, BioLegend), anti-CD90.2 AlexaFluor700 (30H12, BioLegend), anti-IgG1 BV421 (A85-1, BD Biosciences), anti-CD138 BV605 (281-2, BD Biosciences), anti-TCRβ BV711 (H57-597, BD Biosciences), anti-B220 BV750 (RA3-6B2, BioLegend), and anti-IgD BV786 (BD Biosciences). Stained cells were washed twice with 10% DMEM and suspended in the same media containing propidium iodide (PI, Sigma-Aldrich). Labeled cells were analyzed/sorted using BD FACSymphony A5 (BD Biosciences) or BD FACSymphony S6 (BD Biosciences). Flow cytometric data were analyzed with FlowJo software (Treestar Inc.). Dead cells and doublets were excluded from analysis based on propidium iodide (Sigma-Aldrich) staining and FSC-A/FSC-H gating, respectively. GC B cells were identified as TCRβ^-^B220^hi^CD138^-^CD38^lo^GL7^+^ and plasma cells were identified as TCRβ^-^B220^-^CD138^hi^.

### Mouse single B cell culture (‘Nojima’ culture)

Mouse GC B cells were expanded in the presence of NB21.2D9 feeder cells as described^37^. Single B cells were directly sorted into separate wells of 96-well plates pre-seeded with NB21.2D9 cells in 200 µl B cell media (BCM): RPMI-1640 (Invitrogen) with 10% FCS (HyClone, Cytiva), 55 µM 2-mercaptoethanol, 10 mM HEPES, 1 mM sodium pyruvate, 100 units/ml penicillin, 100 μg/ml streptomycin, and non-essential amino acids (Invitrogen) supplemented with recombinant murine IL-4 (2 ng/ml; Peprotech). Cultures were maintained at 37°C with 5% CO2. After two days of culture, 100 µl of culture medium was removed and replaced with 200 µl of fresh BCM. On culture days 2, 3, 5, 6, 7 and 8, 200 µl of culture medium was replaced with fresh BCM. On culture day 10, culture supernatants were harvested for screening the reactivity of secreted clonal IgGs. Culture plates containing GC B cell clones were stored at −80°C until use for V(D)J amplification.

### ELISA

Presence of IgG in culture supernatants from individual wells was determined by ELISA. High-binding 384-well microplates were coated with 2 µg/ml each anti-mouse Igκ and Igλ capture antibodies in 0.1 M sodium carbonate buffer (pH 9) overnight at 4°C. After washing with PBS plus 0.1% Tween 20, the plates were blocked with PBS plus 0.5% BSA at RT for 1 hr. Culture supernatants were diluted 1:10 with PBS containing 0.5% BSA and 0.1% Tween 20. Diluted supernatants (50 μl) were added to the plates and incubated for 2 h at RT, or overnight at 4°C. After extensive washing, HRP-conjugated goat anti-mouse IgG secondary Ab (1:5000 in PBS with 0.5% BSA and 0.1% Tween 20; Southern Biotech) was added and bound HRP activity visualized using a TMB substrate kit (BioLegend). Background signal at 650 nm was subtracted from the signal at 450 nm to calculate the OD_450_ on a Spectramax plate reader (Molecular Devices).

### Multiplex (Luminex) bead assay

Antigen-specific IgG from serum samples and individual culture supernatants were determined in a Luminex assay (Luminex Corp.) as described (Kuraoka et al., 2016). Culture supernatants or sera were diluted 1:10 in Luminex assay buffer (PBS plus 1% BSA, 0.05% NaN3 and 0.05% Tween 20) plus 1% non-fat milk and incubated for 2 h at RT (or overnight at 4°C). Sera were further diluted in 3-fold serial dilutions. Dilutions were then added to a mixture of antigen-coupled microsphere beads in 96-well filter-bottom plates (Millipore). After washing, beads were incubated at RT for 1 h (or overnight at 4°C) with PE-conjugated goat anti-mouse IgG (Southern Biotech). After washing, beads were suspended in assay buffer and analyzed on a Bio-Plex 3D Suspension Array System (Bio-Rad). The following antigens were coupled with carboxylated beads (Luminex Corp): goat anti-mouse Igκ and goat anti-mouse Igλ (both Southern Biotech), negative controls (OVA, rPA, insulin, BSA, streptavidin) native CH848, 10.17DT, ferritin nanoparticle, and heterologous HIV antigens (BG505 SOSIP and CH505 SOSIP). For each IgG^+^ culture supernatant sample and serum sample, concentrations of 10.17DT-binding IgG and total IgG were determined in reference a titration curve of the monoclonal DH270.6 bnAb. AvIn is the ratio of IgG_10.17DT_/IgG_total_ of each sample^38^.

### Ab V(D)J rearrangement amplification and analysis

Rearranged V(D)J gene sequences for mouse GC cells from single-cell cultures were obtained by RT-PCR in one of two methods, both previously described^32; 37^. In both methods, total RNA was extracted with the Quick-RNA 96 Kit (Zymo Research) and treated with DNase I. For cDNA synthesis, two options were used: (1) Superscript III reverse transcriptase with oligo(dT)20 primers, or (2) SMARTScribe Reverse Transcriptase (Clontech) together with 0.2 μM of gene-specific reverse primers and a 1 μM 5′ SMART template-switching oligo containing plate-specific barcodes. The resulting cDNA was amplified by two rounds of semi-nested PCR with Herculase II fusion DNA polymerase (Agilent Technologies) using combinations of forward and reverse primers. Amplicons generated with Superscript III were gel-purified and submitted to the Duke DNA Sequencing Facility for Sanger sequencing. Amplicons generated with SMARTScribe were barcoded, pooled, purified, and sequenced by DNA Link Inc. or CD Genomics Inc. using the PacBio SMRT platform. For each V(D)J sequence, the expected number of sequencing errors was calculated from Phred-scaled base quality scores using custom scripts, and sequences with more than two expected errors were excluded from downstream analyses.

### Recombinant IgG expression and purification

DNA encoding H- or L-chain variable domains was cloned into expression vectors harboring the constant regions of mouse or human IgG1, Igk, Igl, or mouse IgG2c. IgGs were produced by transient transfection of Expi293F cells with the Expifectamine 293 transfection kit (Thermo Fisher), according to the manufacturer’s instructions. Five days post-transfection, supernatants were harvested, clarified by low-speed centrifugation, mixed 1:1 with Protein G binding buffer (for mouse IgG1) or Protein A binding buffer (for human IgG1 or mouse IgG2c), and incubated overnight with Pierce Protein G or Protein A agarose resin (Thermo Fisher). The resin was collected in a chromatography column, washed with binding buffer, eluted in Pierce IgG Elution Buffer (Thermo Fisher), neutralized by 1M Tris (pH 9), and dialyzed into PBS. IgG concentrations were determined with a NanoDrop spectrophotometer (Thermo Fisher).

### HIV neutralization assay

Env-pseudotyped virus neutralization assays completed were measured as a function of reductions in luciferase (Luc) reporter gene expression after a single round of infection in TZM-bl cells ^39; 40^. TZM-bl cells (also called JC57BL-13) were obtained from the NIH AIDS Research and Reference Reagent Program, as contributed by John Kappes and Xiaoyun Wu. Briefly, a pre-titrated dose of virus was incubated with serial dilutions of antibodies in duplicate for 1 h at 37 °C in 96-well flat-bottom culture plates, followed by addition of freshly trypsinized cells. One set of control wells received cells + virus (virus control) and another set received cells only (background control). After 48-72 h of incubation, cells were lysed and measured for luminescence using the Britelite Luminescence Reporter Gene Assay System (PerkinElmer Life Sciences). IC50 and ID80 neutralization titers are the concentration at which relative luminescence units (RLU) were reduced by 50% or 80% compared to virus control wells after subtraction of background RLUs from cell controls.

### Single cell RNA-seq

Following fluorescence-activated cell sorting (FACS), single cells were encapsulated on the Chromium X instrument (10x Genomics, Pleasanton, CA) using the 5′ GEM-X chemistry according to the manufacturer’s protocol. Amplified cDNA was used to generate gene expression (GEX), B-cell receptor (BCR), and Feature Barcode libraries. Libraries were assessed for size distribution using the Agilent TapeStation High Sensitivity D5000 assay (Agilent Technologies, Santa Clara, CA) and quantified using the Qubit fluorometric assay (Thermo Fisher Scientific, Waltham, MA). Pooled libraries were sequenced on the Illumina NextSeq 2000 platform (Illumina, San Diego, CA) using the recommended read configuration (28 × 10 × 10 × 90 cycles).

### Single cell RNA-seq Computational Analysis

Sequencing data from single-cell experiments were processed using the CellRanger software suite (10x Genomics) to perform alignment to a custom mouse genome, barcode processing, and gene expression quantification. For immune repertoire analysis, BCR sequences were assembled and quantified using the corresponding Cell Ranger VDJ pipeline. Downstream analyses were performed in R using Seurat^41^ and custom scripts. Cells were filtered based on standard quality control metrics, followed by normalization, identification of highly variable genes, and data integration. Principal component analysis was performed on the scaled expression matrix, and a shared nearest-neighbor graph was constructed using the selected principal components.

Clustering was conducted using a graph-based Louvain algorithm. Low-dimensional visualization of the resulting clusters was generated using a UMAP projection. Differentially expressed marker genes were used to annotate cluster identities, as described in the main text. Functional, paired BCR sequences were aligned to the DH270 UCA to determine the presence and combination of the four critical mutations.

### Gene Set Enrichment Analysis

Gene set enrichment analysis (GSEA) was performed to identify biological pathways differentially enriched between ipsilateral and contralateral cluster 11 T cells. Differential expression analysis was first conducted at the single-cell level, and genes were ranked by average log₂ fold change (avg_log2FC) between conditions. All expressed genes were included in the ranked list without pre-filtering based on statistical significance. GSEA was carried out using the gseGO function from the clusterProfiler package (4.10.1) in R (4.5.2), querying Gene Ontology Biological Process (GO BP) gene sets. Mouse gene symbols were used as identifiers and mapped internally to the org.Mm.eg.db annotation database. Gene sets containing fewer than 15 genes or more than 500 genes were excluded. Enrichment scores were calculated using a weighted Kolmogorov–Smirnov–like statistic, and normalized enrichment scores (NES) were derived to account for differences in gene set size. Statistical significance was assessed using permutation testing (5,000 permutations), and nominal p-values were adjusted for multiple hypothesis testing using the Benjamini–Hochberg FDR procedure. To allow full inspection of enrichment patterns, all GO BP terms were returned regardless of significance threshold, with statistical significance defined as FDR < 0.05. Enrichment curves and summary dot plots were generated using the enrichplot package.

### Statistics

Statistical significance (p < 0.05) was determined by Wilcoxon matched-pairs signed rank test or Kruskal-Wallis test with Dunn’s multiple comparisons using GraphPad Prism software (version 10.5.0, GraphPad Software). Statistical test is indicated within each figure legend.

## Results

### Generation of F_1_ (DH270 UCA KI x S1pr2^ERT2Cre^-tdTomato) mice to trace the humoral response to an HIV-1 vaccine antigen

To create an experimental model for the analysis of humoral responses to candidate HIV-1 Env vaccine immunogens, we crossed mice homozygous for the V3-glycan DH270 bnAb UCA BCR knock-in (KI) alleles^15^ with the S1pr2^ERT2Cre^-Rosa26^lox-stop-lox-tdTomato^ (S1pr2-RFP) mouse line^36^. The DH270 UCA BCR represents the germline founder of the DH270 bnAb lineage^38^ specific for the V3 loop of the HIV-1 Env. S1pr2-RFP mice were generated in the laboratory of T. Kurosaki^36^ and have been proven useful in tracking the fates of GC lymphocytes^32; 36; 42^. In mice heterozygous for the DH270 UCA V_H_ and V_L_ chains, about 12% of splenic follicular B cells express the DH270 BCR and in their resting state, these cells displayed lower surface densities of membrane IgD^11; 15^. In S1pr2-RFP mice, approximately 70%-80% of early (days 6-8) primary GC B cells and their progeny are permanently marked by tdTomato expression (RFP^+^) following Tamoxifen administration on day 6 post-immunization^32; 36; 42^. The RFP^-^ GC population may include a small number of unlabeled early immigrant GC B cells in addition to a significantly larger population of later (post day 8) GC entrants.

### Characterization of (DH270xS1pr2-RFP) humoral responses to the 10.17DT

(DH270xS1pr2-RFP) mice (**Figure 1A**) were immunized with the CH848 10.17DT stabilized Env trimer (SOSIP) (10.17DT) conjugated to a ferritin nanoparticle (10.17DT-NP). The 10.17DT-NP antigen efficiently activates B cells carrying the DH270 UCA BCR^13; 15^. (DH270xS1pr2-RFP) mice were immunized in the right footpad with 10.17DT-NP in alum. Six days after immunization, tamoxifen was administered *per os*. Serum Ab titers and GC B cell responses were assessed in cohorts of 2–23 mice on days 12, 24, 36, 56-70, and 64-78 after primary immunization (**Figure 1A**), with the overlapping late time points reflecting cohorts sampled before and at a fixed interval after the time of boosting in parallel cohorts.

**Figure 1.**
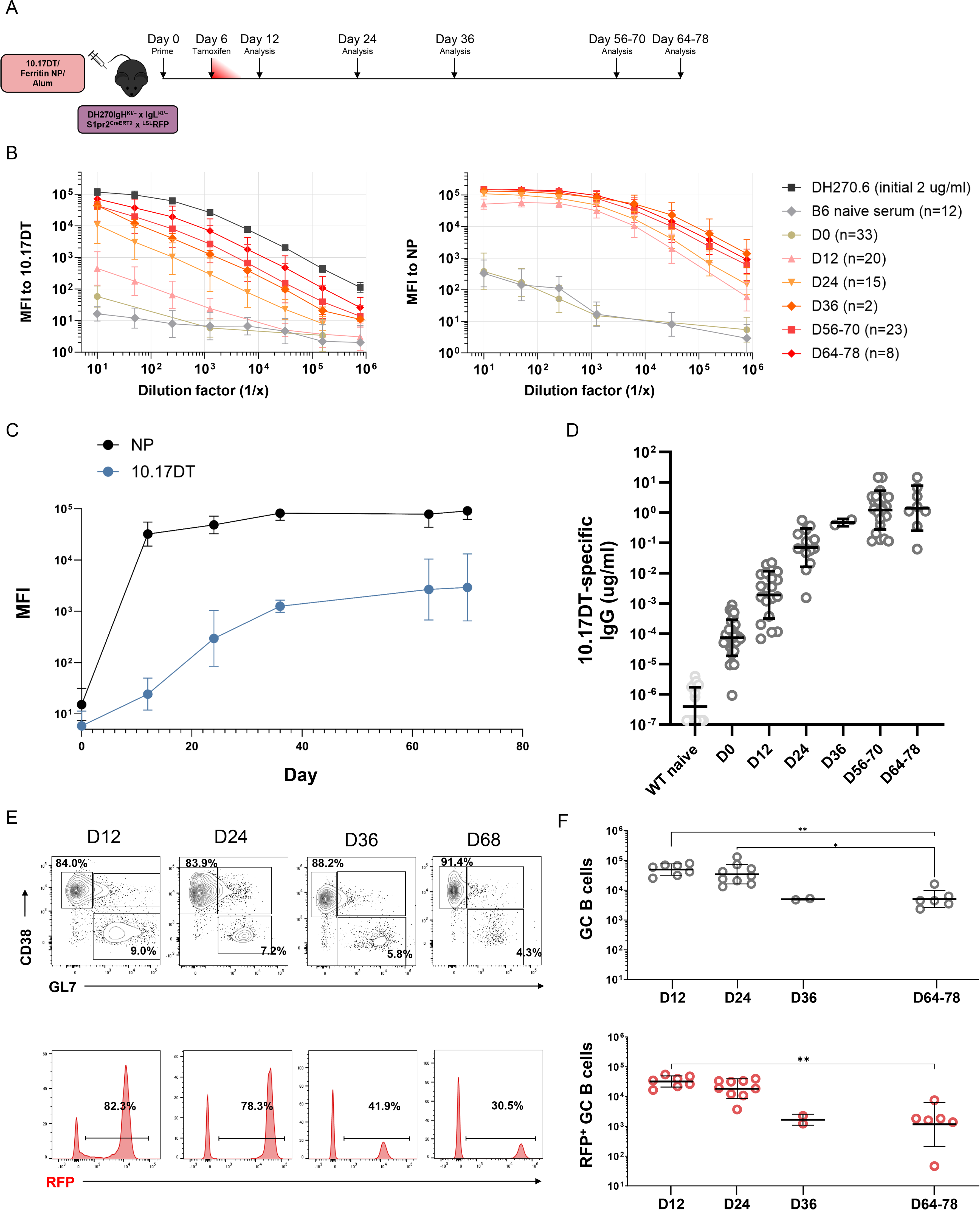
Immunization with 10.17DT-NP elicits persistent humoral responses. **(A)** Experiment schematic. (DH270xS1pr2-RFP) mice were immunized with 10.17DT-NP with alum adjuvant in the right footpad. Tamoxifen was administered on day 6. Analyses were performed at the indicated timepoints. **(B)** Serially diluted serum samples from naïve BL/6 mice and from naïve or immunized (DH270×S1pr2-RFP) mice were assessed individually for binding to 10.17DT (left) or ferritin nanoparticle (right) by Luminex assay; results are displayed as grouped data points by day. Serial dilutions of the mature DH270 lineage antibody, DH270.6 (in black), starting at 2 μg/ml was used as a positive control for 10.17DT binding. Samples at overlapping time points (D56-70 and D64-78) are from separate immunization cohorts. **(C)** Mean fluorescence intensity (MFI) for serum from immunized (DH270×S1pr2-RFP) mice at a 1:1250 dilution, measured over time. Data for D56-70 is plotted at D64 and data from D64-D78 is plotted at D70. **(D)** Concentration of 10.17DT-specific IgG over time following primary immunization. Symbols represent individual mice. Numbers of mice as in Figure 1B. For **(B-D)**, bars represent geometric mean ± geometric SD. **(E)** Representative flow cytometry plots (top row) show the frequency of naïve (B220⁺CD38⁺GL7⁻) and germinal center (GC; B220⁺CD38⁻GL7⁺) B cells, and the proportion of RFP⁺ cells among GC B cells (bottom row). **(F)** Total number of GC B cells (top, grey) and RFP^+^ GC B cells (bottom, red) per lymph node across conditions: D12 (n=7), D24 (n=9), D36 (n=2), and D64-78 (n=6). Bars represent geometric mean ± geometric SD. **, *p* < 0.01; *, *p* < 0.05; by Kruskal-Wallis test with Dunn’s multiple comparisons.

### Serum IgG antibody kinetics after 10.17DT-NP immunization

Serum IgG specific for 10.17DT or NP was quantified in Luminex assays from samples collected between days 0 and 78. Immunization with 10.17DT-NP elicited serum IgG Ab specific for 10.17DT and NP (**Figure 1B-C**). Baseline IgG binding to 10.17DT in unimmunized (DH270 × S1pr2-RFP) mice was higher than in naïve C57BL/6 controls, reflecting low basal DH270 UCA activation (**Figure 1B**). Ab responses to the two vaccine components, 10.17DT and NP, displayed distinct kinetics: whereas 10.17DT-specific IgG increased gradually; the NP response rose rapidly after primary immunization. 10.17DT IgG Ab increased steadily through day 36 and then plateaued over 10 weeks of observation (**Figure 1C-D)**. The concurrent response to the ferritin NP reached near maximal values by day 12 and remained stable over the next 58 days (**Figure 1C**). The measured increase of serum IgG Ab for 10.17DT while unusual for many protein antigens^37; 43; 44^ matches the observed kinetics for other Env antigens^45^. The slow rate at which serum IgG Ab to 10.17DT increased after immunization was not due to the rarity of B cells expressing the DH270 UCA BCR.

### 10.17DT-NP immunization elicits robust and persistent GC responses

Immunization with the 10.17DT-NP vaccine elicited robust and persistent GC responses as determined by flow cytometry. By D12, almost 9% of B220^+^ lymphocytes in the draining popliteal lymph node (LN) had a GC phenotype (B220^+^CD38^lo^GL7^+^), corresponding to a peak of ∼8 x 10^4^ total GC B cells (**Figure 1E-F**). By D36, GC B cell numbers fell significantly (∼4 x 10^3^ cells) but remained stable thereafter (**Figure 1F**). Proportions of IgG1^+^ GC B cells increased progressively from day 12 to day 78, whereas IgM expression was lost completely by D24 (**Figure S1A-B**). Thus, while primary GC responses contracted over time, a reduced but stable population of active GC B cells persisted for 11 weeks.

As described^36; 42^, tamoxifen gavage on day 6 resulted in efficient RFP labeling of GC B cells; 66% on D12 and 58% on D24 postimmunization (**Figure S1C**). With time, RFP^+^ GC B cell frequencies declined, but by D36, reached a stable value of ∼35% that was maintained through D78 (**Figure S1C**). The 10.17DT-NP antigen elicits persistent GC responses that support a stable population of RFP^+^ GC B cells and identifies a persistent equilibrium between early (RFP^+^) and late (RFP^-^) B-cell entrants into primary GCs. This stable equilibrium limits the extent of clonal replacement^23^. 10.17DT-NP immunization establishes a durable GC compartment with stable representation of early-entry, RFP^+^ clones for ≥10 weeks.

### Persistent GCs support ongoing affinity maturation

To determine specificity and avidity of individual GCB cells elicited in the primary response, we cultured single GC B cells in ‘Nojima’ cultures^37^ at D12, D24, and D64-78. After culture, IgG^+^ culture supernatants were screened by Luminex for binding to 10.17DT and NP, as well as the heterologous BG505 and CH505 SOSIPs, and irrelevant proteins as negative controls (**Figure S1D**). Avidity indices (AvIn) were then calculated for GC B cells binding 10.17DT using the mature DH270.6 bnAb as a reference^37^. The DH270.6 bnAb has a dissociation constant (K_D_) of 0.62 nM for 10.17DT^13^ (AvIn = 1.0) and the DH270 UCA Ab, by comparison, has a K_D_ of 532 nM^15^ and an AvIn value of 0.02.

On D12 after immunization, only a small fraction of GC B cell clones was specific for 10.17DT (**Figure 2A**). Between D12 and D24, however, the frequency of 10.17DT-binding cells increased sharply to 68.2% of RFP^+^ and 44.8% of RFP^-^ cells. This frequency continued to rise among RFP^+^ GC B cells to 85.5%, whereas RFP^-^ cells plateaued at ∼36%. Tamoxifen concentrations fall below effective levels by D9, therefore increasing RFP^+^ GC B-cell populations between D12 and D24 must represent local proliferation, not continued immigration^42^. Frequencies of GC B cells specific for the ferritin NP never grew beyond 15% of the GC B-cell population (**Figure S2A**).

**Figure 2.**
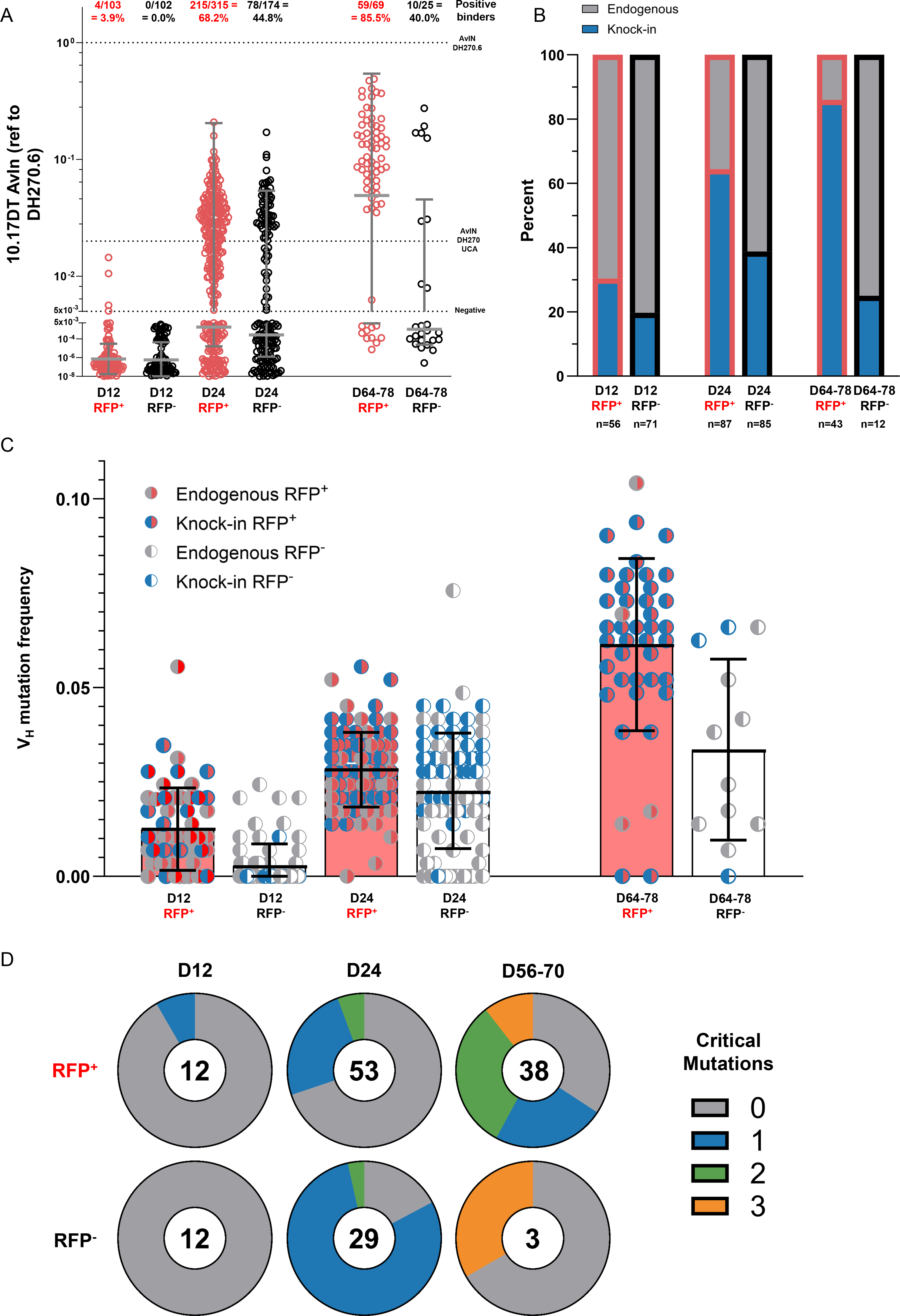
Early-entrant knock-in B cells undergo affinity maturation and acquire hallmark DH270-lineage mutations. (A) Single-cell ‘Nojima’ cultures established from GC B cells from draining lymph nodes taken from a subset mice shown in Figure 1. Analyzed cultures are from multiple mice: D12 (n=2), D24 (n=6), and D64-78 (n=6). Avidity indices (AvIn) referenced to DH270.6 and percentage of cultures with binding (AvIn > 0.005) to 10.17DT. Each symbol represents an IgG⁺ single B cell culture from draining lymph nodes; RFP⁺ cells are shown in red and RFP⁻ cells in black. Dotted lines indicate the negative threshold, the avidity index of the DH270 UCA, and the avidity index of the mature DH270.6 antibody. Bars represent the geometric mean ± geometric SD. (B) Representative cultures underwent BCR sequencing. Cultures for which both heavy and light chain rearrangements were recovered are shown. “Endogenous” includes cells with human–mouse hybrid or fully mouse heavy/light chain pairs. (C) V_H_ mutation frequency among sequenced cultures. Bars represent mean ± SD. Only cells with paired heavy and light chains and heavy-chain V(D)J sequences were included. (D) Pie charts showing the proportion of sequenced wells containing the indicated number (0–4) of critical DH270-lineage mutations. Analysis includes cells with paired KI heavy and light chains.

On D64-D78, 8.7% (6/69) of RFP^+^ GC B cells showed no measurable binding for either antigen. AvIn values for 10.17DT, like 10.17DT-binding frequency, rose markedly between D12 and D24 and increased further by D64–78. On D24, only about 1% (4/315) RFP⁺ cells displayed robust affinity maturation (AvIn >0.1) while 50% (34/70) of RFP^+^ GC B cells did by D64-D78 (**Figure 2A**). The progressive accumulation of both antigen-binding frequency and avidity over months underscores that prolonged GC residency drives the maturation of founder clones.

To understand better the recruitment, persistence, and dynamics of affinity maturation of B cells bearing the DH270 KI BCR, we sequenced V(D)J gene rearrangements from representative subsets of GC B cells from each sample time. Even by D12, early immigrant RFP^+^ GC B cells were more likely to express the DH270 KI BCR (**Figure 2B**). Whereas the proportion of RFP^+^ GC B cells expressing the KI BCR continuously increased between D12 and D64-78 to a peak of 86%, KI BCR expression in the later arriving RFP^-^ population never exceeded 40% of the RFP^-^GC B population (**Figure 2B**). The incomplete enrichment of the DH270 KI BCR within the RFP^−^ population likely reflects competition with established RFP^+^ GC clones that have already undergone affinity maturation, limiting the recruitment and expansion of newly arriving KI-expressing B cells.

To compare the extent of somatic evolution in RFP^+^ and later arriving RFP^-^ GC B cells, we determined V_H_ and V_L_ mutation frequencies of both KI and endogenous BCR gene rearrangements. On D12 the mean frequency of V_H_ mutations in RFP^+^ GC B cells (1.3%) was higher than that of concurrent RFP^-^ cells (0.3%) (**Figure 2C**). The low frequency of mutations in RFP^-^ presumably reflects their later entry into GCs, after tamoxifen labeling. With time, KI V_H_ and V_L_ mutation frequencies increased in both populations, but RFP^-^ GC B cells consistently exhibited lower mutation frequencies than observed in contemporary RFP^+^ samples (**Figure 2C and S2B**). In addition to establishing that RFP^-^ cells reach a stable equilibrium with the founder-derived RFP^+^ population, these data show that a substantial fraction (≈40%) of RFP^-^ cells are antigen-specific and have accumulated somatic mutations, indicating that late GC entrants participate in ongoing selection and diversification. Recovery of two RFP^+^ GC B cells expressing the unmutated DH270 KI BCR on D64-78 is evidence either for re-entry of B cells that exited primary GCs before fixing V(D)J mutations or the long retention of the GC phenotype absent continuing hypermutation (**Figure 2C and S2B**).

Four critical and improbable replacements (G57R and R98T in the heavy chain and S27Y and L48Y in the light chain) in the DH270 UCA BCR provide nearly 80% of the bnAb activity present in the mature DH270 bnAb^13^. These critical mutations in RFP^+^ GC B cells increased over time and, by D64-78, 64.1% of GC B cells in ‘Nojima’ cultures contained at least one of the four critical mutations; 10.3% of GC B cells carried three of the critical mutations necessary for breadth and potency (**Figure 2D**).

### Asymmetric responses to ipsilateral and contralateral homologous boosts

We and others have shown that the location of boosting can profoundly influence the recruitment of antigen-experienced B cells into secondary GC responses^32; 33^ and we asked how this might affect the trajectory of antibody evolution in our model. Thus, we boosted (DH270 × S1pr2-RFP) mice primed 56-70 days earlier with the homologous 10.17DT-NP immunogen in either the right hock (ipsilateral) or in the distal left hock (contralateral), including the D64-78 primary response cohort as a “no-boost” control group (**Figure 3A**).

**Figure 3.**
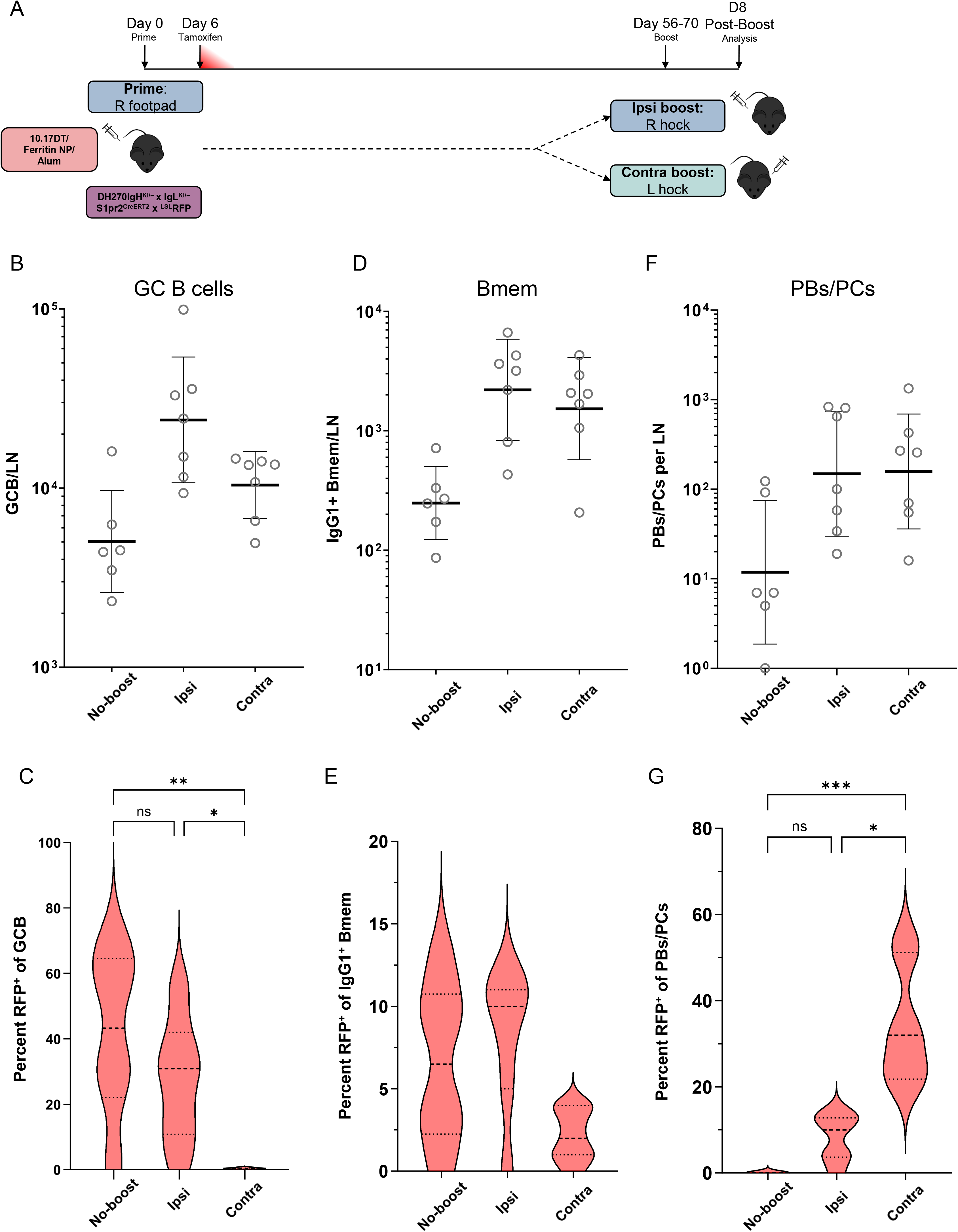
Ipsilateral boost more efficiently recruits progeny of GC B cells that participated in the primary response. **(A)** Experiment schematic. (DH270xS1pr2-RFP) mice were immunized with 10.17DT-NP with alum adjuvant in the right footpad. Tamoxifen was administered on day 6. Boost immunization was administered on days 56-70 either at the right hock (ipsilateral boost) or left hock (contralateral boost) and analysis was performed eight days later. Analyses represent the draining popliteal lymph node after boost unless otherwise indicated. **(B)** Number of B220^+^CD138^−^GL-7^+^CD38^lo^ GC B cells in the draining LNs of no-boost (n=6), ipsilateral (n=7), or contralateral boost (n=7) and the left, non-draining LN without boost (n=3) and after ipsilateral boost (n=2). Circles represent individual mice and bars represent geometric mean with geometric SD. **(C)** Percentage of RFP^+^ cells of GC B cells from draining lymph nodes. Horizontal lines represent first quartile, median, and third quartile. **(D)** Number of IgD^-^IgG1^+^ Bmem from the same LNs as **(C)**. Geometric mean with geometric SD. **(E)** Percentage of RFP+ cells of IgD^-^IgG1^+^ Bmem. Horizontal lines represent first quartile, median, and third quartile. **(F)** Number of B220^-^CD138^hi^ PBs/PCs from the same LNs as **(C)**. Geometric mean with geometric SD. **(G)** Percentage of RFP^+^ cells of PBs/PCs. Horizontal lines represent first quartile, median, and third quartile. ***, *p* < 0.01; **, *p* < 0.01; *, *p* < 0.05; ns > 0.05; by Kruskal-Wallis test with Dunn’s multiple comparisons.

Eight days after boosting, GC B cell numbers in the boosted ipsilateral LN increased five-fold over “no-boost” controls, to an average of 3.2 x 10^4^ B cells. Approximately one-third of these GC B cells were RFP^+^ (**Figure 3B, S3A**). Although contralateral boosts also elicited robust GCs (mean=1.1 x 10^4^ GC B cells; **Figure 3B, S3A**) these GCs contained almost no RFP^+^ GC B cells (**Figure 3C**). Numbers of RFP^+^ GC B cells in the ipsilateral LN were >220-fold higher than in the contralateral node.

Because of the downregulation of IgD on naïve B cells in DH270 KI mice^15^, we used class-switched IgD^-^ IgG1^+^ to identify *bona fide* Bmem. Bmem populations expanded comparably at both sites, from ∼3×10³ cells in no-boost controls to ∼3×10⁴ ipsilateral and ∼2×10⁴ contralateral (**Figure 3D**). Unlike GC B cells, RFP+ Bmem were detectable at the contralateral site (2.5%), though at roughly one-fourth the frequency seen at the ipsilateral site and in no-boost controls (∼10%) (**Figure 3E**). RFP^+^ Bmem therefore circulate to distal sites but fail to seed productive GCs there.

Local plasmablast (PB)/plasmacyte (PC) responses also differed between ipsilateral and contralateral boosts as described previously^33^. Although PB/PC numbers also increased comparably after both boosts (**Figure 3F, S3B**), the RFP^+^ fraction was markedly higher after contralateral boosting (∼30%) than ipsilateral (∼10%) (**Figure 3G**). Despite this asymmetry in RFP^+^ PB/PC differentiation, both ipsilateral and contralateral boosts elicited comparable, ∼5-fold, increases in serum Ab (**Figure S3C-D**). These observations demonstrate a strong bias for RFP^+^ B cells to fuel post-boost, ipsilateral GC responses but to favor PB/PC differentiation after contralateral boosts. Distal boosting promotes PB/PC differentiation in the progeny of primary GC B cells, whereas local boosts lead to GC B-cell engagement. These observations are the first confirmation of GC and PB/PC asymmetry recently reported by Dhenni et al.^33^

### Delayed homologous boosting advances critical mutation accumulation and elicits neutralization breadth

In prior studies, closely spaced, repeat immunizations of DH270 mice with 10.17DT-NP did not produce B cells carrying all four critical mutations^13; 15^. We therefore asked whether delayed re-exposure to antigen could better advance maturation of these evolving lineages. We characterized post-boost B-cell repertoires with single-cell ‘Nojima’ cultures of GC B cells recovered eight days after ipsilateral (n=473) or contralateral boosts (n=418)^37^. After ipsilateral boosts, RFP^+^ GC B cell populations comprised some 85% binding to 10.17DT and 7% specific for NP (**Figure 4A and S4A**), values essentially identical to that of contemporary no-boost controls (**Figures 2A and S2A**). AvIn values for pre-boost and ipsilateral post-boost RFP^+^ GC B cells were also similar, with distributions of AvIn measures ranging from 0.05 to 0.50 (5% - 50% of the DH270.6 bnAb). Whereas DH270 V_H_ mutation frequencies were also comparable in the no-boost and ipsilateral boost groups (**Figure 4B-4C**), ipsilateral boosting substantially increased the proportions of RFP^+^ GC B cells carrying mutations critical for neutralization breadth^20^, with 26% of GC B cells harboring three critical mutations (**Figure 4D**).

**Figure 4.**
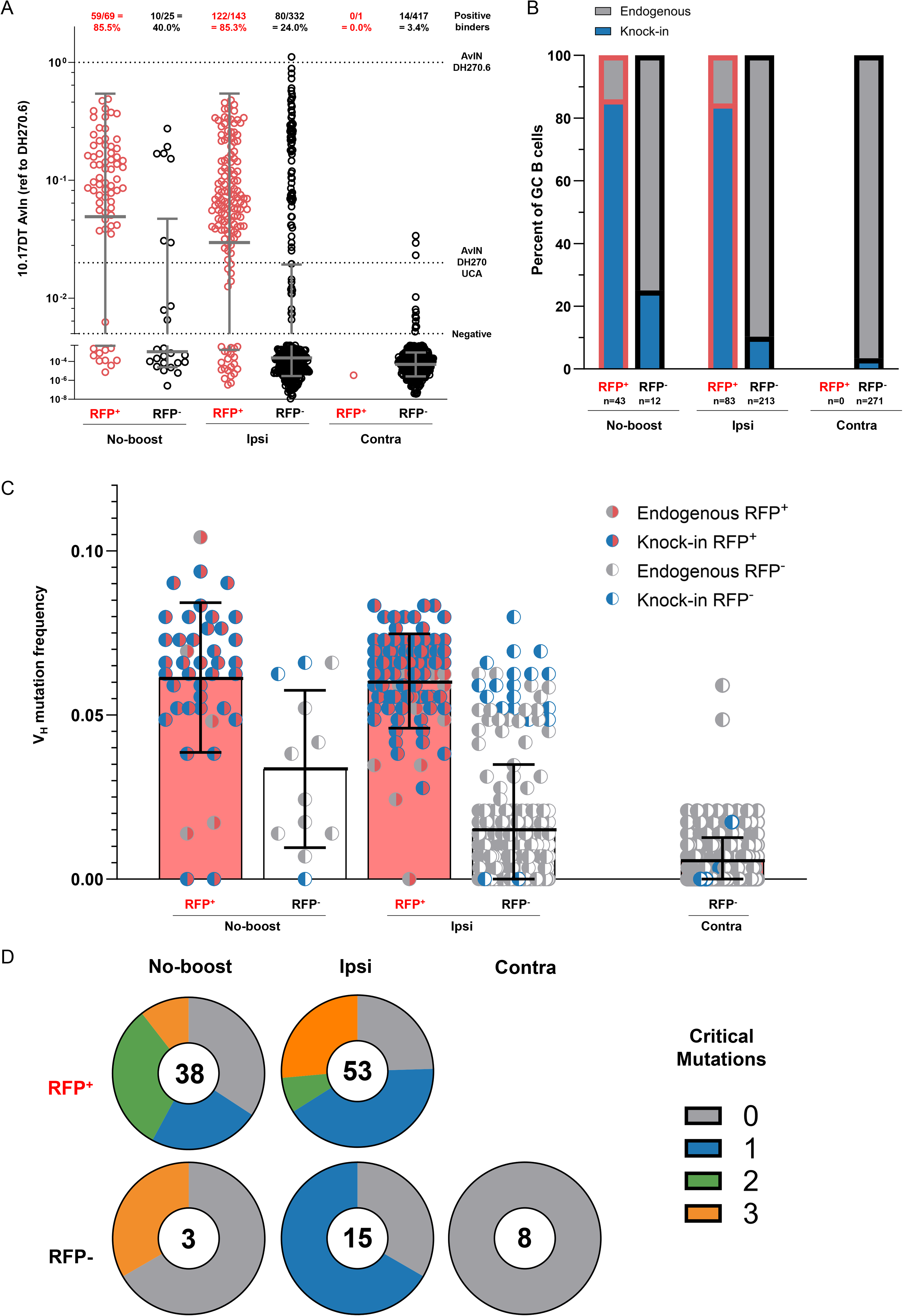
Ipsilateral boost expands high-avidity clones with combinations of critical mutations. (A) Single-cell ‘Nojima’ cultures were performed from draining lymph nodes of mice analyzed in Figure 3. Cells are pooled from multiple mice: no-boost (n=6), ipsilateral (n=7), and contralateral (n=7). Avidity indices and percentage of cultures with binding to 10.17DT (AvIn > 0.005). Each symbol represents an IgG⁺ single B cell culture from draining lymph nodes; RFP⁺ cells are shown in red and RFP⁻ cells in black. Dotted lines indicate the negative threshold (AvIn = 0.005), the avidity index of the DH270 UCA, and the avidity index of the mature DH270.6 antibody. Bars represent the geometric mean ± geometric SD. (B) Representative subsets of cultures underwent BCR sequencing. Shown are cultures from which both heavy and light chains were recovered. No sequenced antibodies were recovered from RFP^+^ cells at contralateral boost sites. “Endogenous” includes cells with human–mouse hybrid or fully mouse heavy/light chain pairs. (C) V_H_ mutation frequency among sequenced cultures. Bars represent mean ± SD. Only cells with paired heavy and light chains and heavy-chain V(D)J sequences. (D) Pie charts showing the proportion of sequenced wells containing the indicated number (0–4) of critical DH270-lineage mutations. Analysis includes cells with knock-in heavy and light chains.

Ipsilateral boosts also expanded RFP^-^ GC B cell numbers (**Figure 3B-C**), the great majority expressing endogenous BCR rearrangements (**Figure 4B**). Nonetheless, clones from within this RFP^-^ GC population exhibited the highest AvIn values for 10.17DT with occasional antibodies binding as avidly as the mature DH270 bnAb (**Figure 4A**). BCR rearrangements from six RFP^-^GC B cell clones with AvIn > 0.5 were fully endogenous; the mouse Ab repertoire is capable of avid response to the 10.17DT SOSIP.

In contrast, contralateral boosts elicited robust de novo GC responses (**Figure 3B**) that inefficiently recruited both RFP^+^ and KI BCR B cells. AvIn values were markedly lower in the contralateral boost group (**Figure 4A**). Only a single RFP^+^ GC B cell was recovered from contralateral boost ‘Nojima’ cultures (0.2%), only 3% expressed the KI BCR, and only 4% bound 10.17DT or NP (**Figures 4A, 4B and S4A**). The infrequent KI BCR GC B cells generally carried low V-region mutation frequencies (**Figure 4C and S4B**). Recruitment of KI BCR B cells to contralateral boost site GCs even fell below that of the 12% frequency of KI BCR among peripheral B cells^13^. In this model system, contralateral boosts disfavor recruitment of both RFP⁺ and RFP^-^ B cells expressing KI BCRs into secondary GC responses, potentially due to Ab feedback^46^.

Asymmetric boost responses were also observed for GC B cells expressing endogenous BCRs specific for NP. While approximately 7% of GC B cells bound NP after ipsilateral boosts, only ∼1% did so after contralateral boosts, and these cells generally exhibited weaker NP binding (**Figure S4A**). Suppression of antigen-specific B-cell recruitment to GCs after contralateral boosting is not an artifact of the elevated precursor frequencies of the DH270 BCR KI.

### Heterologous binding and neutralization breadth

Beyond the acquisition of mutations and of avidity to 10.17DT, we wanted to examine the development of cross-reactivity to heterologous HIV antigens and neutralization breadth. Analysis of the specificity and avidity of IgG secreted by clonal ‘Nojima’ cultures from both primary and secondary GC responses revealed distinct binding patterns for 10.17DT, NP, and the two heterologous SOSIPs BG505 and CH505, with no evidence for off-target binding or polyreactivity (**Figure 5A-B**). RFP⁺ cell cultures were enriched both for avid 10.17DT binding and cross-reactivity with heterologous HIV Env products, consistent with somatic evolution leading to reactive breadth as well as increasing affinity. To determine the basis of this breadth, we identified clonal cultures that secreted IgG reactive to both the 10.17DT and BG505 SOSIPs. Of 50 cultures with detectable binding to both the 10.17DT and BG505 SOSIP, 44 (88%) were from either no-boost (18, D64-78) or ipsilateral boost (26) cohorts (**Figure 5C**).

**Figure 5.**
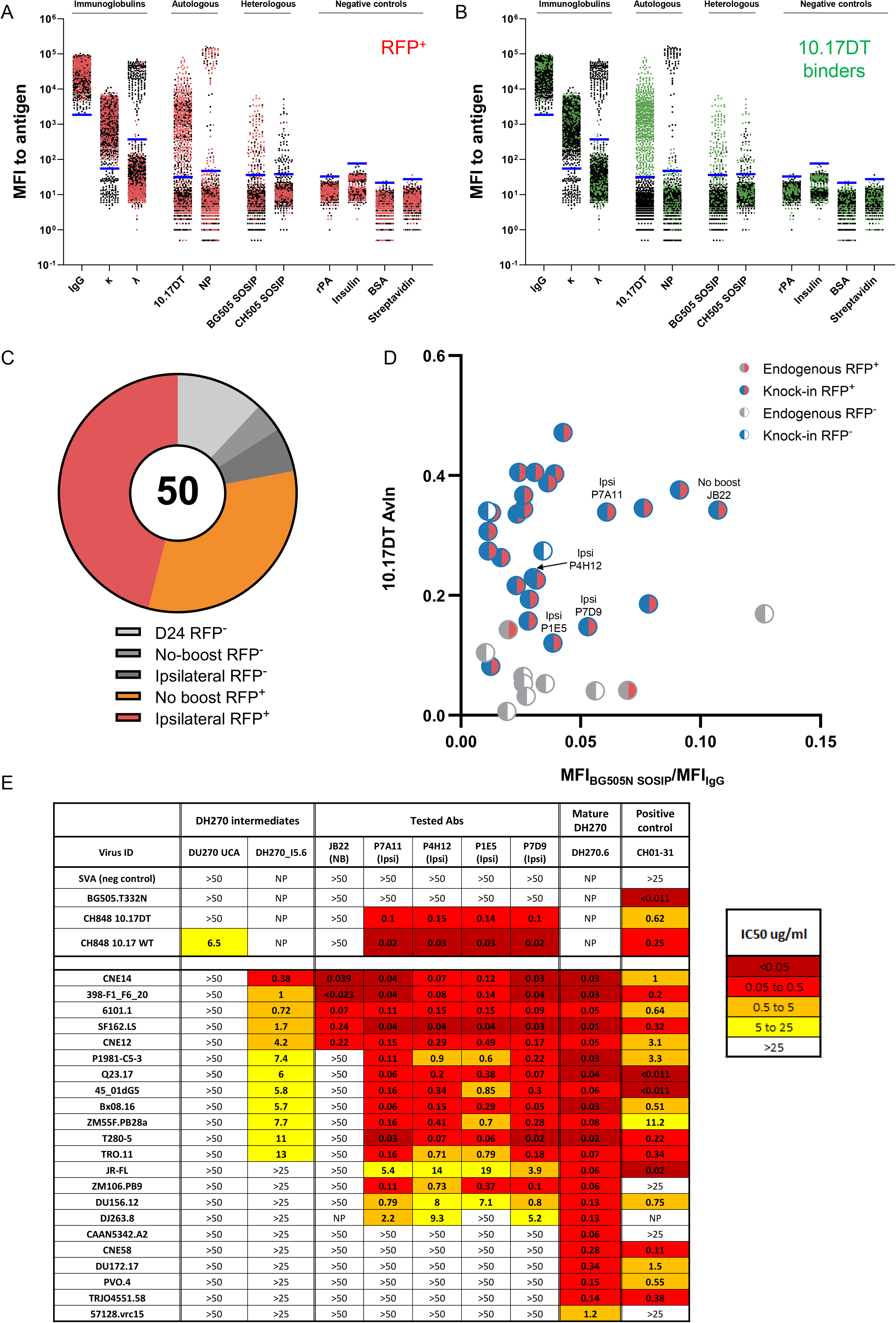
‘Nojima’ cultures reveal development of HIV cross-reactivity in early-entrant GC B cells. **(A-B)** Mean fluorescence intensity (MFI) of IgG in ‘Nojima’ culture supernatants binding to antigen-conjugated beads in a Luminex assay for immunoglobulins, autologous antigens, and negative controls including cultures after primary and secondary immunization (n=1,739). Subsets of supernatants were tested against heterologous antigens (BG505 SOSIP n=1,261, CH505 SOSIP n=1,487). Each symbol represents clonal IgG derived from a single GC B cell. Horizontal blue lines indicate the binding threshold (mean + 6 SD of signal from control wells containing no B cells). (**A)** RFP^+^ cells are highlighted in red (**B)** 10.17DT binding-supernatants are highlighted in green. **(C)** Fifty clones with detectable binding to both 10.17DT and BG505N were identified as cross-reactive. Pie chart shows the distribution of these cross-reactive clones by experimental condition. **(D)** BCR rearrangements were sequenced from 37 of the 50 cross-reactive clones. Clones with detectable BG505N binding that underwent BCR sequencing are displayed, colored by BCR type and RFP status. **(E)** Neutralization capacity of select cross-reactive antibodies cloned from single-cell ‘Nojima’ cultures tested against viruses selected for sensitivity to DH270.6 and heterologous HIV-1 viruses. NP = not performed, NB = No-boost, Ipsi = ipsilateral boost

Thirty-seven paired heavy- and light- chain sequences were recovered from this cross-reactive group, of which 27 expressed the DH270 KI BCR (**Figure 5D**). Five cross-reactive clones taken from different mice and spanning a range of avidities were selected for functional virus neutralization testing (**Figure 5D**). Neutralization was assessed against wild-type CH848 10.17, autologous 10.17DT, and BG505.T332N, and a panel of 22 heterologous viruses sensitive to DH270.6 neutralization^13^ (**Figure 5E**). Four of five tested KI BCR clones displayed neutralization breadth falling between the DH270 lineage intermediate I5.6 and the mature DH270.6 bnAb^13^; all four derived from ipsilateral boosts, while the fifth, JB22, a high avidity clone recovered from a no-boost cohort, showed minimal breadth. Significant neutralization breadth, nearly 75% of the mature DH270 bnAb, can develop in early secondary GC responses with long intervals between prime and boost immunizations.

### scRNA-seq reveals acquisition of four critical mutations and disfavored mutational pathways

‘Nojima’ cultures established that mutation accumulation advances progressively during the primary response and further after delayed ipsilateral boosting in GC B cells, but cannot resolve which whether GC-egressed populations carry mutations on productive trajectories toward bnAb activity. To address these questions, and to identify the early cellular events underlying the GC and PC asymmetry observed at day 8 after boosting (**Figure 3**), we performed single-cell transcriptomics on RFP^+^ lymphocytes isolated from draining LNs. Cells were isolated at 36 hours after boosting to capture differences in the cells first responding to antigen at each site before antigen-driven proliferation obscures the composition of the GC.

We isolated and pooled B220^+^RFP^+^ and CD3^+^CD4^+^RFP^+^ cells from the draining LNs of mice after ipsilateral (n=14) or contralateral boosts (n=23) performed 10, 13, or 19 weeks after primary immunization as well as from contemporary no-boost controls (n=18). RFP^+^ LN cells from mice 16 days after primary immunization (n=2) were included to facilitate transcriptomic identification of B- and T-cell differentiation types (**Figure 6A and Figure S5A**). Flow cytometric enumeration, showed about 600 RFP^+^ B cells per LN at 10-19 weeks after priming (**Figure 6B**). By 36 hrs after ipsilateral boost, numbers of local RFP^+^ B cells increased >3-fold to an average of some 2,000 cells per LN (**Figures 6B**). Contralateral boosts resulted in much lower numbers of RFP^+^ B cells, with average numbers <100 cells in boosted contralateral LNs (**Figure 6B**).

**Figure 6.**
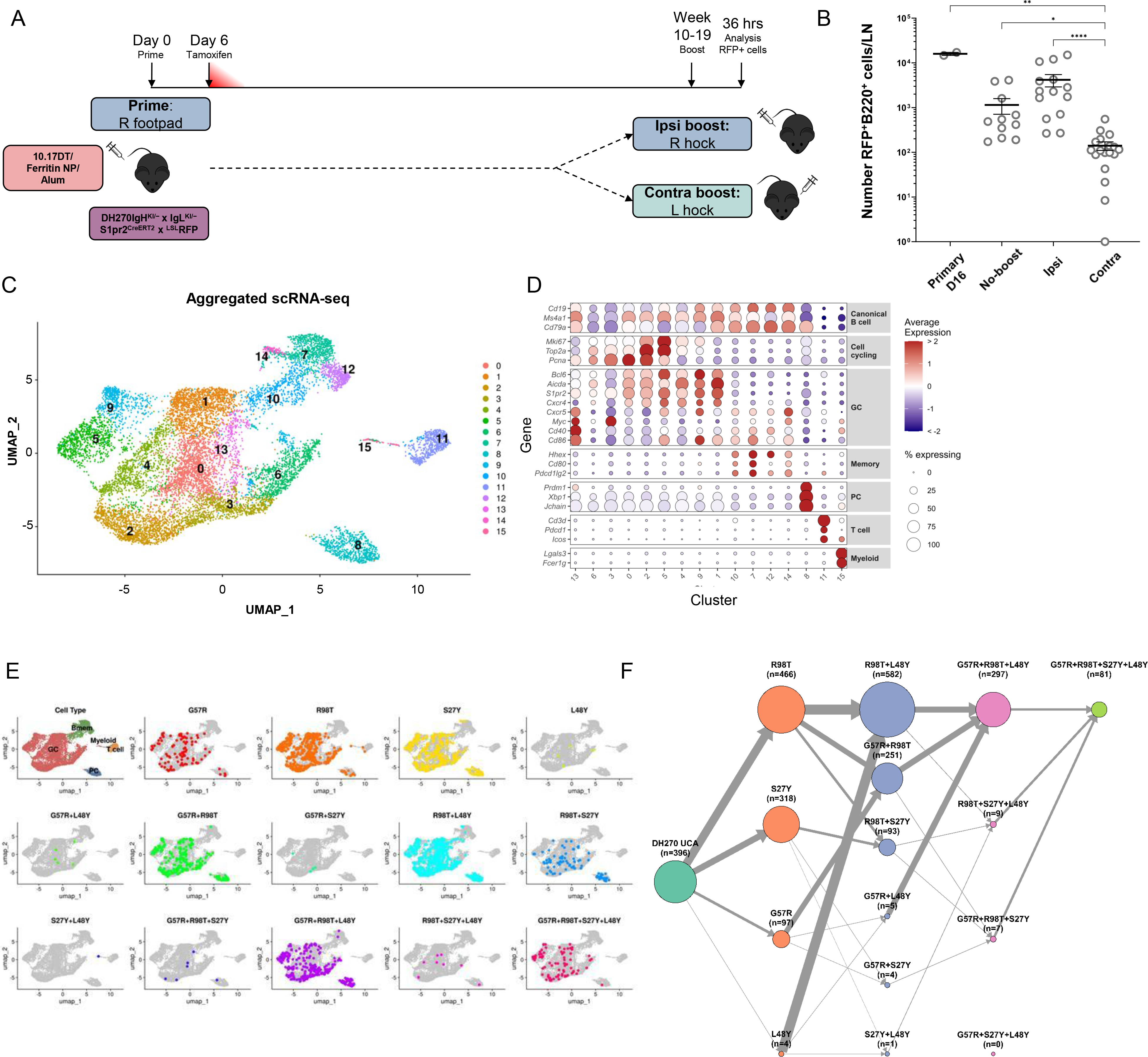
scRNA-seq reveals disfavored mutational pathways. **(A)** Experiment schematic. (DH270xS1pr2-RFP) mice were immunized with 10.17DT-NP with alum adjuvant in the right footpad. Tamoxifen was administered on day 6. Single-cell RNA sequencing of lymph node RFP^+^ B (B220^+^) and CD4^+^ T cells following primary immunization, no-boost, ipsilateral boost, or contralateral boost at the indicated time points. **(B)** Total number of RFP⁺B220⁺ cells per lymph node. Horizontal bars represent the arithmetic mean ± SEM. ****, *p* < 0.0001; **, *p* < 0.01; *, *p* < 0.05; Kruskal–Wallis test with Dunn’s multiple-comparisons correction. **(C)** Uniform manifold approximation and projection (UMAP) of all sequenced cells (n=12,526) pooled from four experimental conditions (D16 primary immunization, no-boost, ipsilateral, contralateral) colored by transcriptional cluster identity. **(D)** Heatmap of scaled gene expression used to define transcriptional clusters. **(E)** Distribution of DH270 UCA–derived clones across transcriptional clusters identified by scRNA-seq, utilizing the same dataset as in Figure 6. Gray points represent all analyzed cells, colored points indicate cells carrying the indicated DH270 lineage mutations or mutation combinations. **(F)** Directed network diagrams showing frequencies of all observed combinations of the four critical mutations derived (G57R, R98T, S27Y, and L48Y) from scRNA-seq. Each node represents a unique mutation combination, with node size proportional to the number of cells observed carrying that genotype. Node color indicates the total number of critical mutations present. Arrows indicate observed transitions between mutation states, with edge thickness proportional to the number of cells in the receiving node.

Unsupervised clustering of the entire scRNA-seq dataset (n=12,526 cells over all four experimental conditions) identified fifteen transcriptionally distinct clusters (**Figure 6C**) which were visualized using a UMAP projection. Nine clusters represented GC B cells, defined by canonical GC markers (*S1pr2*, *Aicda*, *Bcl6*) in combination with *Cxcr4* (dark zone), *Cd86* (light zone), and cell cycle markers (*Myc* and *Mki67*)^47–49^ (**Figure 6D**). Three GC clusters (13, -6, -3) contain light-zone (LZ) or selected LZ states; four clusters (0, -2, -5, -4) comprise dark-zone (DZ) B cells in distinct cell-cycle phases; and two (9, -1) identify DZ-to-LZ or LZ-to-DZ transition states. Four clusters (10, -7, -12, -14) exhibited Bmem transcriptional programs, including upregulation of *Hhex*, *Pdcd1lg2*, and *Cd80*. The remaining three clusters contained PBs/PCs (cluster 8; *Prdm1*, *Xbp1*, *Jchain*), T cells (cluster 11; *Cd3d*), and a small group of ill-defined myeloid cells (cluster 15; *Lgals3*, *Fcer1g*).

We next examined the distribution of mutations critical for DH270 neutralization breadth (G57R, R98T, S27Y, and L48Y)^13^ across B-cell populations. RFP^+^ B cells containing one or more critical mutations were identified throughout GC populations and among PBs/PCs, but were much less frequent within memory B-cell clusters (**Figure 6E**). GC cells with all four critical mutations were identified and were recovered exclusively following ipsilateral boosting.

Consistent with continued evolution over prolonged periods, mutation frequencies among ipsilateral boost GC B cells continued to increase through week 19 after primary immunization (**Figure S5B**). Indeed, 97% of cells carrying all four critical mutations were recovered at the 19-week time point, indicating that prolonged GC activity continues to drive accumulation of breadth-associated mutations long after immunization.

To determine whether accumulation of all four critical mutations conferred the expected functional properties in the context of naturally evolved V(D)J repertoires, we selected five clonal IgG Abs encoded by KI BCR rearrangements (JB1-JB6) containing all four critical mutations and tested for neutralization breadth in comparison to the DH270 UCA, the I5.6 lineage intermediate, and the mature DH270 bnAb. Four (JB1, JB3-6) of these five IgG Abs showed increased neutralization breadth compared to the I5.6 intermediate, approaching breadth and potency of the DH270 bnAb (**Figure S5C**).

Comparison of the frequencies of B cells with BCR containing ≥1 critical mutation regardless of boosting condition identified favored and disfavored evolutionary pathways leading towards bnAb activity (**Figure 6E**). Critically, the order in which critical mutations accumulate varied significantly within this small set of mutations. For example, while BCRs with the R98T+L48Y paired mutations were most abundant in our sample (n=582), that pair of mutations most likely followed an earlier, primary R98T substitution (n=466) versus L48Y (n=4) (**Figure 6F**).

Similarly, whereas a primary S27Y substitution was frequent (n=318), early acquisition of S27Y was an evolutionary dead-end for DH270 bnAb maturation. Mutational pairs (G57R, n=4; L48Y, n=1) and triplets (G57R+R98T, n=7; R98T+L48Y, n=9) containing S27Y were rare. The mutation pair S27Y+R98T (n=93) likely represents a fitness barrier preventing further maturation of a DH270 bnAb following immunization with 10.17<u>DT</u> (**Figure 6F**).

### Early determinants of boost site asymmetry

While site-dependent asymmetry in secondary humoral response is well established, the basis of the asymmetry is poorly understood. A central question is whether ipsilateral boosting drives superior GC responses simply by expanding the persistent GC population already resident in the draining LN, or whether it additionally recruits circulating antigen-experienced memory B cells into new GC responses. More broadly, the early cellular populations that are recruited at each site and seed the divergent outcomes observed have not been well defined. We used the 36-hour time point to distinguish between these possibilities by characterizing the composition and mutational states of RFP^+^ cells first recruited at each site.

Despite the extended interval after primary immunization, the transcriptional organization of persistent GC responses remained remarkably stable. The nine GC clusters identified in late primary responses occupied the same light-zone, dark-zone, and transitional states observed during active primary responses, despite being separated by 8–17 weeks (**Figures 7A-B**). These findings indicate that long-lived GCs elicited by 10.17DT-NP remain transcriptionally active and retain canonical GC architecture months after immunization, connecting to the increased mutations over time.

**Figure 7.**
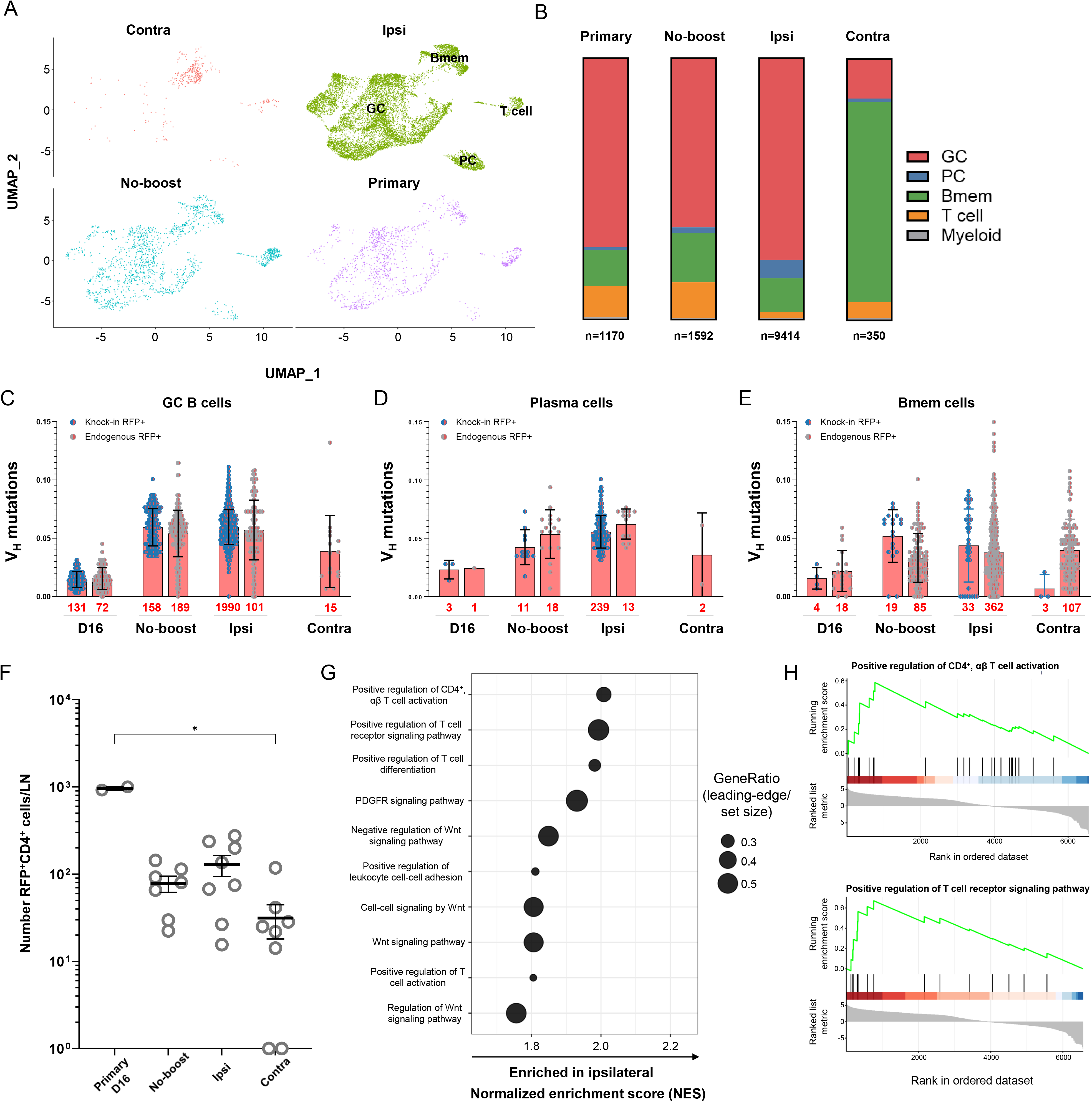
Boost sites are composed of distinct B and T cell subsets. **(A)** UMAP colored by experimental condition, showing distribution of cells across clusters following primary immunization (n=1,170), no-boost (n=1,592), ipsilateral boost (n=9,414), or contralateral boost (n=350). **(B)** Frequency of clusters grouped by transcriptional phenotype and experimental condition **(C-E)** V_H_ mutation frequency by experimental condition within GC **(C)**, plasma cell **(D)**, and memory B cell **(E)** compartments, comparing knock-in RFP⁺ and endogenous RFP⁻ cells. Total number of cells within each category under bars. **(F)** Total number of RFP⁺CD4⁺ cells per lymph node. Horizontal bars represent the arithmetic mean ± SEM. *, *p* < 0.05; Kruskal–Wallis test with Dunn’s multiple-comparisons correction. **(G)** GSEA was performed using genes ordered according to average log₂ fold change between ipsilateral and contralateral cluster 11 T cells. The top ten Gene Ontology (GO) Biological Process terms ranked by normalized enrichment score (NES) are shown. Positive NES values indicate enrichment toward ipsilateral cells. **(H)** Running enrichment score plots for the GO Biological Process terms *positive regulation of CD4⁺ αβ T cell activation* (top) and *positive regulation of T cell receptor signaling pathway* (bottom). Genes are ranked by average log₂ fold change (ipsilateral versus contralateral cluster 11 T cells), with positive values indicating higher expression in ipsilateral cells. Vertical black ticks indicate the positions of genes belonging to each pathway within the ranked list, and the green line represents the running enrichment score. Leftward enrichment reflects preferential representation among ipsilateral-enriched genes.

Following ipsilateral boosting, the number of RFP^+^ GC B cells increased markedly (**Figure 6B**) along with substantial increases in the ratios of GC B cells (1:1 to 20:1) and plasmacytes (1:1 to 18:1) expressing the KI BCR to endogenous BCRs (**Figure 7C-D**). In contrast, the KI ratio within the memory B-cell compartment shifted in the opposite direction, from approximately 1:4 in no-boost controls to 1:11 after ipsilateral boosting (**Figure 7E**). The reciprocal depletion of KI-expressing Bmem and enrichment of KI-expressing GC cells is consistent with rapid recruitment of antigen-experienced DH270-lineage memory cells into secondary GC responses (**Figure 7C**). Close examination of the changes in V_H_ mutation frequencies across conditions further support the presence of memory-cell recruitment. First, the distribution of V_H_ mutation frequencies among KI-expressing GC B cells broadened substantially following ipsilateral boosting (**Figure 7C**). Second, 40 of 1,990 KI-expressing GC B cells recovered after ipsilateral boosting possessed mutation frequencies below the lowest value observed in no-boost controls (**Figure 7C**). The appearance of these less-mutated cells within the post-boost GC compartment is only explained by influx of circulating antigen-experienced cells rather than expansion of resident GC populations. Early plasmacytes elicited by ipsilateral boosts had V_H_ mutation frequencies less disperse than GC B cells and uniformly increased compared to contemporary no-boost controls (**Figure 6D**). These early, post-boost plasmacytes likely represent selective, affinity-driven, plasmacyte differentiation from Bmem immigrants^50^.

In contrast, contralateral boosts recruited a fundamentally different population of antigen-experienced cells. Bmem clusters dominated the small (n=<100; **Figure 6B**) RFP^+^ B-cell populations recovered at 36 hours after contralateral boosts, representing 77% of recovered RFP^+^ cells (**Figure 7A-B**). The predominance of RFP^+^ Bmem at 36 hours after contralateral boosts, coupled with the subsequent enrichment of RFP^+^ plasmacytes and absence of RFP^+^ GC B cells at D8 (**Figure 3D,G**), suggests that the differentiation of recalled RFP^+^ memory cells is restricted to the plasmacytic fate. Despite being readily detectable, RFP^+^ Bmem BCRs were encoded virtually only by endogenous gene segments, although they were mutated at frequencies comparable to RFP^+^ KI cells recovered after ipsilateral boosts (97%, **Figure 7E**). Thus, contralateral boosting did not efficiently recruit the DH270 lineage that dominates persistent GC responses and gives rise to broadly neutralizing activity. Instead, it preferentially engaged a distinct endogenous memory B-cell population with unknown specificity.

Given the importance of T-cell help in shaping humoral responses^51; 52^, we also examined RFP^+^CD4^+^ T cells recovered from all control and experimental boost groups. S1pr2-Cre is active in GC-resident Tfh cells^42^, and the RFP^+^CD4^+^ cell clusters retained high expression of canonical Tfh markers, including *Pdcd1* and *Icos* (**Figure 6D**). Prior work has demonstrated a critical role for antigen-specific memory Tfh cells in the magnitude of Bmem reactivation^33^. Low numbers,

≈100 of RFP^+^ CD4^+^ T cells were present in both no-boost control and ipsilateral boost animals; values that represent a 10-fold reduction compared to D16 primary responses (**Figure 7F**). RFP^+^ CD4 T cell numbers were even less frequent (10-fold reduction) in contralateral boost LNs (**Figure 7F**). Gene set enrichment analysis (GSEA) of ranked differential expression between ipsilateral and contralateral cluster 11 T cells revealed consistent trends toward enrichment of T-cell activation related programs in ipsilateral cells, including positive regulation of T-cell receptor signaling, CD4⁺ T-cell activation, and T-cell differentiation (normalized enrichment score [NES] ∼1.8–2.0). None of these pathways, however, reached statistical significance after correction for multiple testing (false discovery rate (FDR) > 0.05) (**Figure 7G-H**).

## Discussion

Using a novel model that combined a lineage-tracing approach with HIV bnAb UCA BCR knock-in mice and single-cell analyses, we show that persistent GCs support continued affinity maturation of founder clones and local, ipsilateral, boosts preferentially engage these lineages in secondary GCs. This model achieved broadly neutralizing DH270-lineage antibodies after a single primary immunization and a single homologous boost. Moreover, by tracking antigen-experienced B cells, we provide the first independent replication of boost-site asymmetry in GC versus PC fate determination and identify novel mechanisms governing this asymmetry, including differences in local T cell help availability and the distinct identities of Bmem populations recruited to ipsilateral versus contralateral sites. Finally, we illustrate preferred and disfavored evolutionary pathways within the DH270 lineage, underscoring an important phenomenon for vaccine strategies.

There is increasing appreciation for the persistence of GCs elicited by immunization of animals^23; 35^ and humans^53; 54^. Strategies to prolong GC duration have been developed, including escalating antigen doses or modulating the timing of immunization^55; 56^. Here, we show that a single dose of 10.17DT-NP immunogen induces GCs that persist for at least three months and persistent RFP^+^ cells remain strikingly similar in overall composition to those present at day 16 after immunization. A small (≈35% GC B cells) but durable (≤19 wks) population of RFP^+^ GC-founder B cells continue to accumulate critical mutations and increasing avidity. Similar long-lived, local populations were observed after influenza and SARS-CoV-2 infection^23^, where naïve “invaders” were proposed to restrict founder-cell proliferation. While the initial decrease in RFP^+^ GC B cells is consistent with naïve invasion, the continued cycling, mutation, and affinity maturation of this persistent founder subset argues against restriction of their proliferation over time. The early-entrant GC population is physiologically significant; long after primary immunization nearly all RFP^+^ founder-derived GC B cells robustly bound antigen and accumulated improbable BCR mutations critical for neutralization breadth and potency^8^. Notably, prior studies using the same knock-in model with multiple serial immunizations did not achieve a comparable degree of acquisition of critical mutations or neutralization breadth^13; 15^, suggesting that allowing prolonged, uninterrupted time in GCs is a critical factor in supporting bnAb development.

For *lineage design* strategies, once B cells acquire mutations critical for bnAb activity, methods to promote their recall or retention in GC responses are necessary for continued maturation. Eight days after ipsilateral boosting, some 35% of GC B cells were derived from the primary GC response whereas very few RFP^+^ GC B cells were detected following contralateral boosting despite similarly sized germinal center responses. After contralateral boosts, even low-affinity unmutated knock-in BCRs were disfavored, comprising only ∼3% of GC B cells, lower than their frequency in the naïve repertoire (∼12%)^15^ or at D12 after primary immunization (∼20-30%). Antibody feedback, whereby antigen-specific circulating Ab modulates GC responses, likely limits the participation of epitope-specific BCR knock-in B cells^46; 57–60^ and unless serum antibody is depleted, secondary GCs are largely composed of naïve cells^25; 51^. The dominance of naïve and non-specific memory B cells in contralateral nodes illustrates this point and while diversification via these naïve lineages may benefit responses to antigenically variable pathogens, it is counterproductive for lineage-based vaccine strategies that depend on iterative maturation of defined B-cell clones. We show here and in our prior work that targeting the same draining lymph nodes through local boosting overcomes this limitation and enables antigen-experienced B cells to participate in secondary GC responses. Importantly, only GC B cells after ipsilateral boost acquired critical mutations associated with neutralization breadth. Ipsilateral boosting thus represents a practical strategy to enhance affinity maturation towards bnAb development.

While ipsilateral boosting promoted GCs containing RFP^+^ B cells, contralateral boosting favored differentiation of RFP⁺ B cells into PCs, replicating findings from Dhenni et al.^33^ in an independent model, suggesting that boost location imposes a bias between the GC and PC fates. We provide two, non-mutually exclusive mechanistic insights to explain this phenomenon. First, the location of antigen encounter may bias B cell fate through *extrinsic* cues from the local microenvironment. Dhenni et al.^33^ demonstrated that primed subcapsular sinus macrophages retained in draining lymph nodes promoted local Bmem retention and GC re-entry and that Tfh depletion prior to boosting reduced GC recall responses. We now show that *early* after contralateral boosting the local lymph nodes contained fewer RFP⁺ CD4^+^ T cells with reduced transcriptional signatures of activation, suggesting that abundant and specific Tfh availability favors GC reformation and disfavors PC differentiation.

Second, intrinsic differences among RFP^+^ B cell populations may influence differentiation fates. After formation, Bmem are generally thought to exit into systemic circulation as “patrolling” cells^50^ and Bmem exported from the primary site may differ in their capacity to form PCs^34^. Dhenni et al.^33^ demonstrated that the Bmem pool present at distal and local boost sites differ. Our data extend this observation: whereas Dhenni and colleagues examined antigen-specific B cells, we analyzed all GC-derived RFP⁺ cells and identified a dominant population of distinct Bmem present early after contralateral boosting. Unlike the RFP^+^ GC B cells that predominate after ipsilateral boost, these cells did not express the DH270 UCA BCR, but rather endogenous rearrangements with extensive somatic hypermutation. Therefore, Bmem present at distal sites are largely not specific for the immunizing antigen and instead may recognize unrelated environmental antigens. While these Bmem are present early after contralateral boosting, they do not go on to form GCs; GC B cells 8 days after contralateral boosts are almost exclusively RFP^-^and unmutated. Therefore, the RFP⁺ B cells recruited to ipsilateral versus contralateral responses are fundamentally different populations, with antigen-specific, avid GC B cells dominating at the local site and extensively mutated, non-specific Bmem at the distal site. These intrinsic differences may themselves impose distinct differentiation fates independent of the microenvironment. Taken together, our findings show that both the local microenvironment in the form of retained, antigen-experienced CD4^+^ T cells and the intrinsic properties of RFP^+^ B cells shape whether secondary responses favor the GC or PC differentiation fates.

An ongoing debate is regarding whether ipsilateral boosting represents “re-fueling” of persistent GCs or secondary GCs. Prior studies have shown that location-dependent recall responses are preserved even after the disruption of organized GCs, with local boosts continuing to preferentially generate GCs enriched for antigen-experienced B cells^32; 33^. Our experiments offer considerable support for secondary GCs. First, only 36 hours after ipsilateral boosts we observe a marked shift, from ∼1:1 to ∼20:1, in the ratio of knock-in to endogenous BCR GC B cells. The magnitude of this change is unlikely to result solely from the proliferation of small persistent GC B-cell populations, especially given that large antigens like 10.17DT-NP require some 8 hours before their detection on follicular dendritic cells^61; 62^. Second, and more definitively, the expanded populations of GC B cells after ipsilateral boosting include cells with fewer V-region mutations than those present in no-boost control GCs. RFP^+^ GC B cells with lower frequencies of BCR mutations cannot represent the products of long persistent GC B cells with higher mutation numbers. Together, these observations are consistent with a substantial contribution to secondary GC responses elicited by ipsilateral boost. Importantly, for lineage design strategies, the distinction is largely irrelevant, as both mechanisms support iterative rounds of selection and mutation within antigen-specific clones.

Beyond boosting strategy, this study also identifies preferred and disfavored evolutionary pathways within the DH270 bnAb lineage. In particular, the S27Y light-chain substitution was rarely observed in combination with other critical mutations, indicating that early acquisition of this mutation may represent an evolutionary dead-end. Although this pattern is specific to the DH270 lineage, it highlights a broader principle relevant to lineage-based vaccine design: successful antibody maturation may depend not only on acquiring particular mutations but also on the order in which they arise. Certain substitutions may alter the structural context of the antigen-binding site in ways that constrain subsequent mutational trajectories. In this model, mutations such as S27Y and L48Y may be necessary for achieving neutralization breadth but are better tolerated later in affinity maturation, after other substitutions establish a permissive structural framework. These findings suggest that some evolutionary pathways toward broadly neutralizing activity are intrinsically disfavored, an important consideration for vaccine strategies that aim to guide antibody lineages through sequential immunization.

## Acknowledgements

We are grateful for the helpful assistance of Masayuki Kuraoka, Dongmei Liao, Xiaoe Liang, Ling Yuan, Xiaoyan Nie, and Scott Szafranski.

## Author contributions

John S. Barber (Conceptualization, Methodology, Formal analysis, Investigation, Data curation, Writing—original draft, Writing—review & editing, Visualization, Funding acquisition); Keisuke Tonouchi (Investigation, Writing—review & editing); Chen-Hao Yeh (Conceptualization, Methodology, Investigation, Writing—review & editing); Madison Berry (Formal analysis, Visualization); Helene F. Kirshner (Formal analysis, Visualization); Kevin Wiehe (Formal analysis, Visualization, Writing—review & editing); Amanda Eaton (Investigation); David C. Montefiori (Investigation); Ming Tian (Resources, Writing—review & editing); Frederick W. Alt (Resources, Writing—review & editing); Kevin O. Saunders (Resources); George M. Shaw (Resources, Writing—review & editing, Funding acquisition); Barton F. Haynes (Conceptualization, Methodology, Resources, Writing—review & editing,

Funding acquisition); Garnett Kelsoe (Conceptualization, Methodology, Resources, Writing—original draft, Supervision, Funding acquisition).

## Abbreviations

AvIn: avidity index
BCM: B cell culture medium
Bmem: memory B cell
bnAb: broadly neutralizing antibody
Contra: contralateral (boost)
DZ: dark zone
Env: HIV-1 envelope
FDR: false discovery rate
GC: germinal center
GO: Gene Ontology
GSEA: gene set enrichment analysis
ID80: 80% inhibitory dose
Ipsi: ipsilateral (boost)
KI: knock-in
LN: lymph node
Luc: luciferase
LZ: light zone
MFI: mean fluorescence intensity
NB: no boost
NES: normalized enrichment score
NP: nanoparticle
PB: plasmablast
PC: plasmacyte
PI: propidium iodide
RFP: red fluorescent protein
RLU: relative luminescence units
scRNA-seq: single-cell RNA sequencing
SOSIP: stabilized (native-like) Env trimer
UCA: unmutated common ancestor
UMAP: uniform manifold approximation and projection
WT: wild-type.

## Supplemental figure titles and legends

**Supplemental figure 1.**
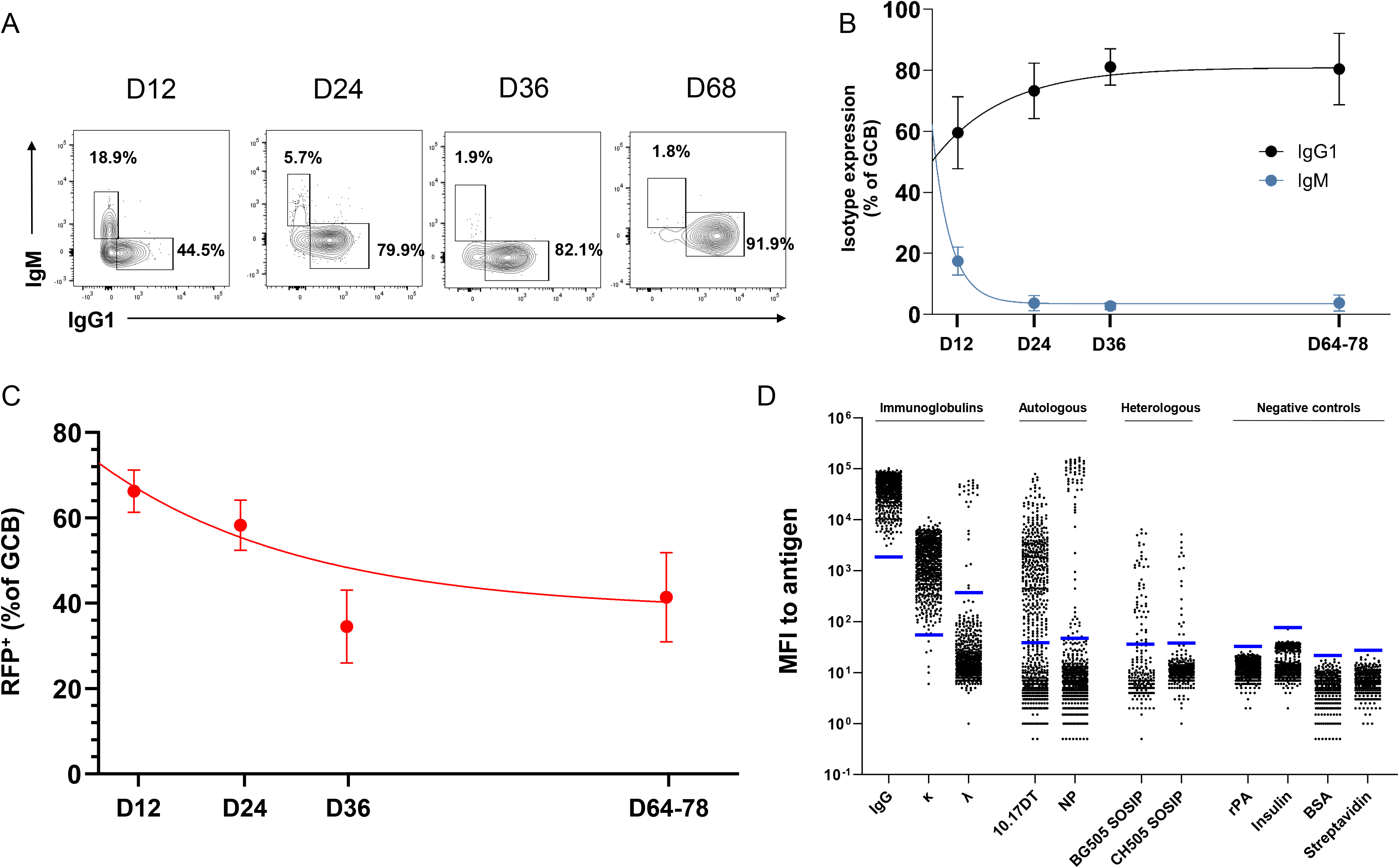
Isotype expression, clonal persistence, and antigen-specificity GC B cells after primary immunization. **(A)** Representative flow cytometry plots showing IgM and IgG1 expression among GC B cells. **(B)** Quantification of IgG1 and IgM expression among GC B cells. Mean ± SD and one-phase decay best-fit curve shown. **(C)** Frequency of RFP^+^ amongst GC B cells at D12 (n=7), D24 (n=9), D36 (n=2), and D64-78 (n=6). Mean ± SD and one-phase decay best-fit curve shown. **(D)** Mean fluorescence intensity (MFI) of IgG in ‘Nojima’ culture supernatants binding to antigen-conjugated beads in a Luminex assay for immunoglobulins, autologous antigens, and negative controls after primary immunization (n=789). Subsets of supernatants were tested against heterologous antigens (BG505 SOSIP n=262, CH505 SOSIP n=488). Each symbol represents clonal IgG derived from a single GC B cell. Horizontal blue lines indicate the binding threshold (mean + 6 SD of signal from control wells containing no B cells).

**Supplemental figure 2.**
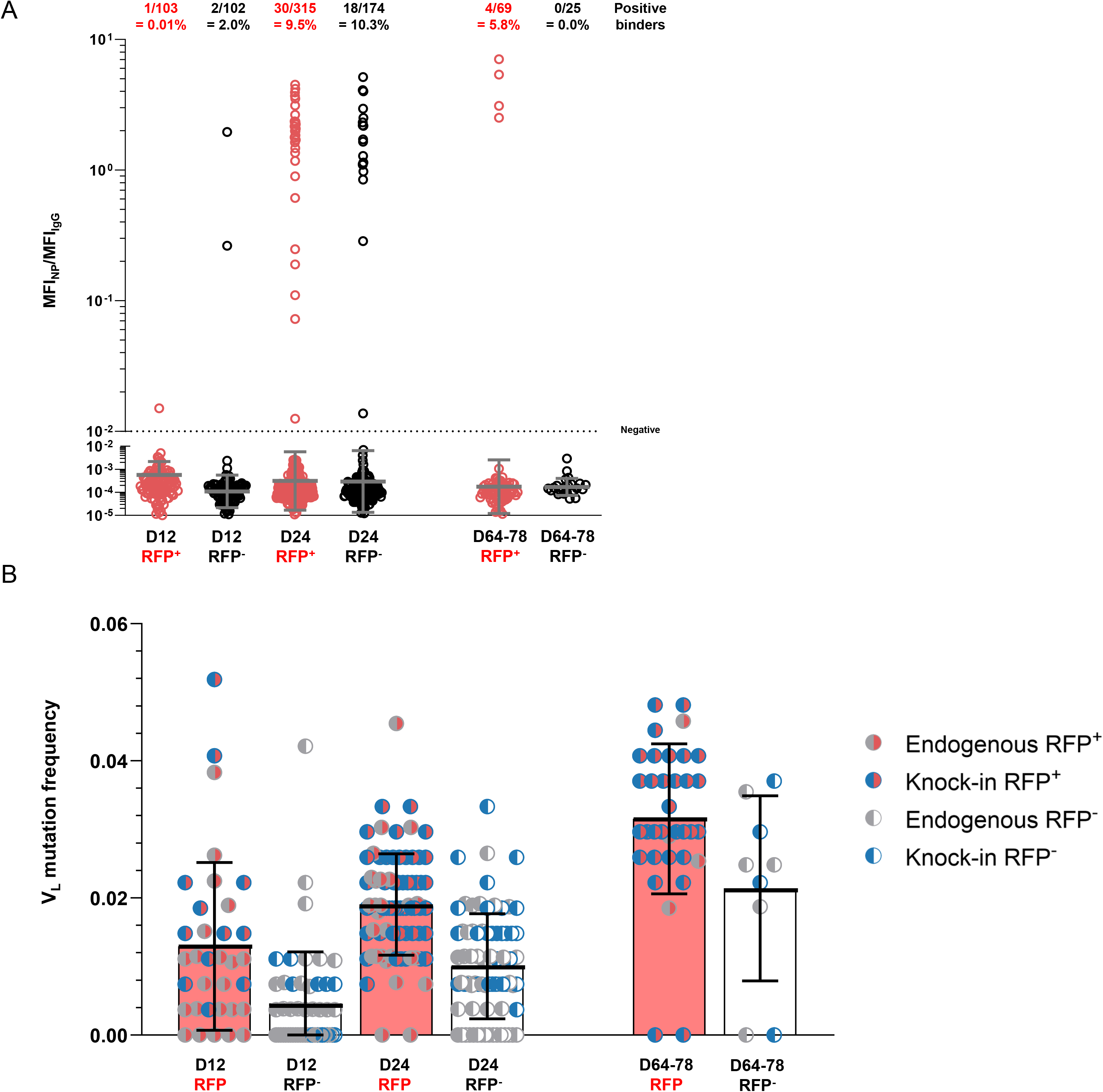
GC B cell reactivity to ferritin nanoparticle scaffold and light chain mutation frequency after primary immunization. (A) Specific activity for ferritin NP of single B cell cultures. Calculated as mean fluorescence intensity (MFI) of ferritin NP divided by MFI of IgG. Above the graph is percentage of cultures with binding to ferritin nanoparticle using a cutoff specific activity of 0.01. Each symbol represents an IgG⁺ single B cell culture from draining lymph nodes; RFP⁺ cells are shown in red and RFP⁻ cells in black. Bars represent the geometric mean ± geometric SD. MFI values of zero for ferritin NP were considered 0.5 (the lowest MFI value of the Luminex assay) for display in log scale and calculation of geometric mean. (B) V_L_ mutation frequency among sequenced cultures. Bars represent mean ± SD.

**Supplemental figure 3.**
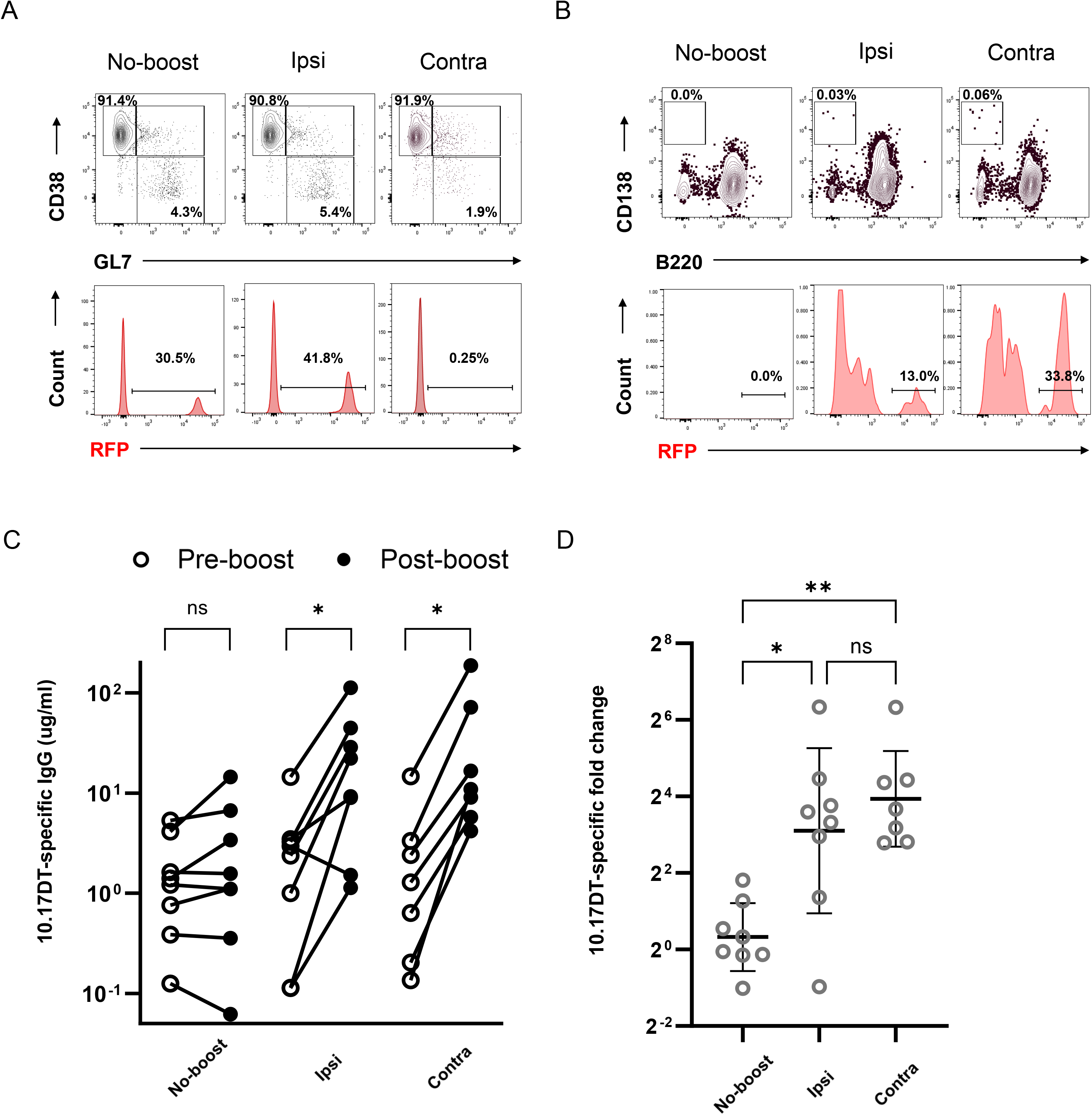
Representative flow cytometry and serologic characterization of mice after boosting. **(A)** Representative flow cytometry plots (top row) show the frequency of naïve (B220⁺CD38⁺GL7⁻) and germinal center (GC; B220⁺CD38⁻GL7⁺) B cells, and the proportion of RFP⁺ cells among GC B cells (bottom row). **(B)** Representative flow cytometry plots (top row) show the frequency of plasmablasts/plasma cells (B220^-^CD138^hi^) B cells, and the proportion of RFP⁺ cells among PBs/PCs (bottom row). **(C)** Concentration of 10.17DT-specific IgG in mice without boost or after ipsilateral or contralateral boost. Lines connect data from the same mouse. *, *p* < 0.05; ns > 0.05; Wilcoxon signed-rank test. **(D)** Fold change in 10.17DT-specific IgG in mice without boost or after ipsilateral or contralateral boost. Symbols represent individual mice; bars indicate the geometric mean and geometric standard deviation. **, *p* < 0.01; *, *p* < 0.05; ns > 0.05; by Kruskal-Wallis test with Dunn’s multiple comparisons.

**Supplemental figure 4.**
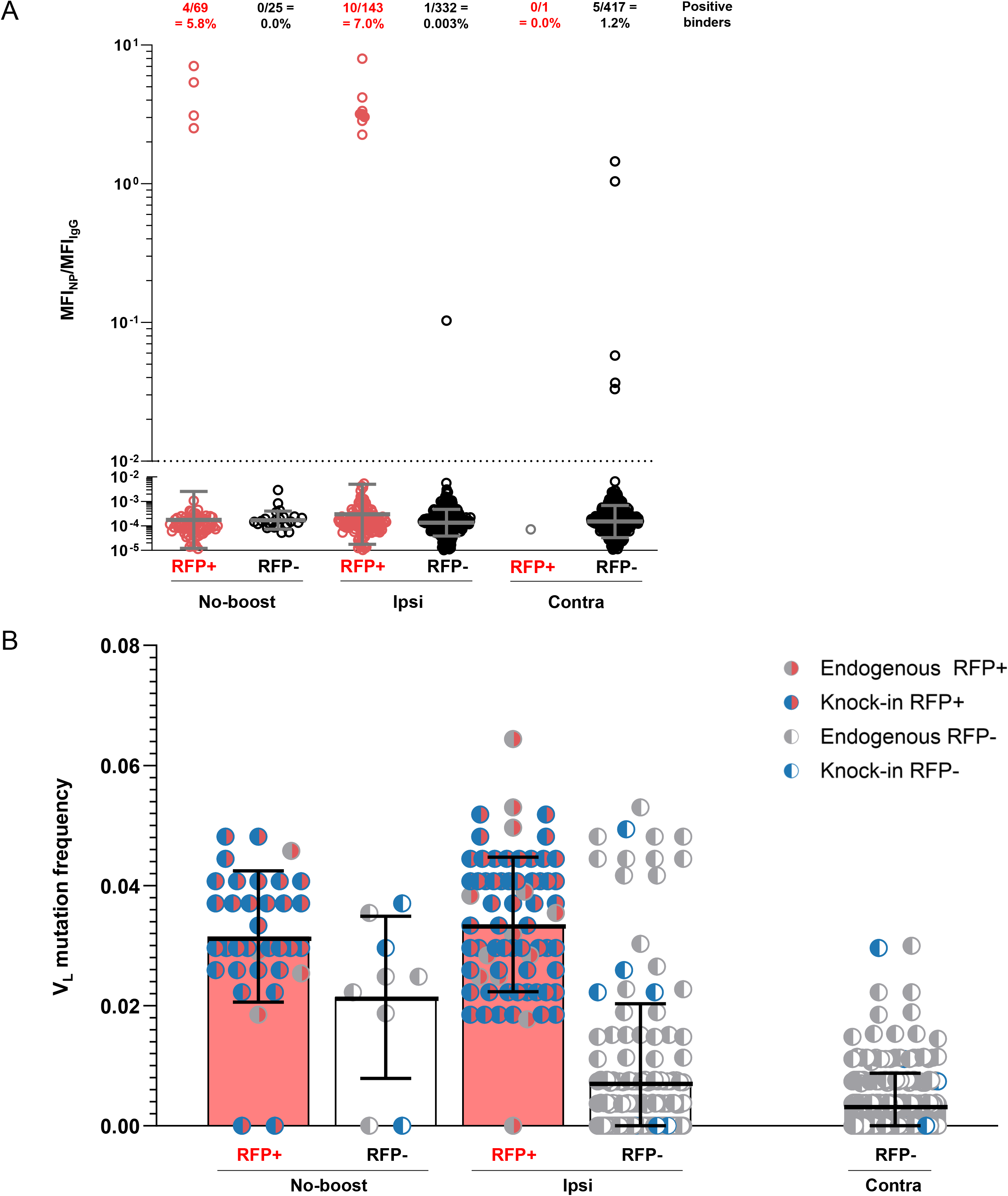
GC B cell reactivity to ferritin nanoparticle scaffold and light chain mutation frequency boosting. (A) Mean fluorescence intensity (MFI) and percentage of cultures with binding to ferritin nanoparticle. Each symbol represents an IgG⁺ single B cell culture from draining lymph nodes; RFP⁺ cells are shown in red and RFP⁻ cells in black. Bars represent the geometric mean ± geometric SD. (B) V_L_ mutation frequency among sequenced cultures. Bars represent mean ± SD.

**Supplemental figure 5.**
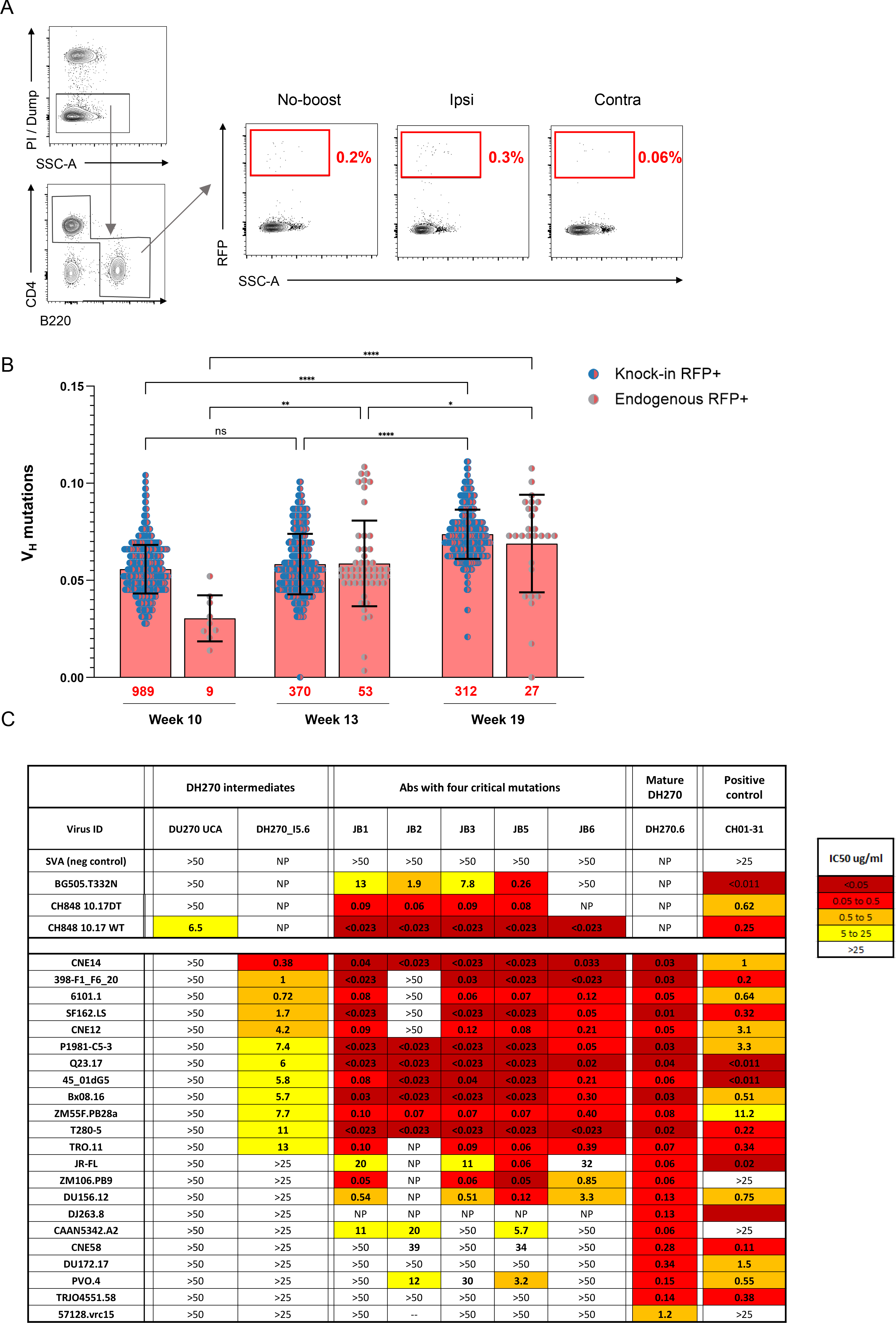
Flow cytometry, BCR sequencing, and neutralization 36 hours after boosting. (A) Representative flow cytometry sorting strategy from an unboosted mouse. Cells were previously gated on single lymphocytes. The dump channel included propidium iodide (PI), CD8α, NK1.1, Gr-1, and F4/80. (B) V_H_ mutation frequency in germinal center (GC) B cells over time after ipsilateral boosting, shown separately for knock-in RFP⁺ (DH270 UCA–derived) and endogenous RFP⁻ populations. (C) Neutralization capacity of antibodies containing all four critical mutations (G57R, R98T, S27Y, and L48Y) cloned from scRNA-seq data and tested against viruses selected for sensitivity to DH270.6 and heterologous HIV-1 viruses

## References

1. Jardine JG, Kulp DW, Havenar-Daughton C, Sarkar A, Briney B, Sok D, Sesterhenn F, Ereno-Orbea J, Kalyuzhniy O, Deresa I, Hu X, Spencer S, Jones M, Georgeson E, Adachi Y, Kubitz M, deCamp AC, Julien JP, Wilson IA, Burton DR, Crotty S, Schief WR. Hiv-1 broadly neutralizing antibody precursor b cells revealed by germline-targeting immunogen. Science. 2016;351(6280):1458–1463. 10.1126/science.aad9195

2. Havenar-Daughton C, Sarkar A, Kulp DW, Toy L, Hu X, Deresa I, Kalyuzhniy O, Kaushik K, Upadhyay AA, Menis S, Landais E, Cao L, Diedrich JK, Kumar S, Schiffner T, Reiss SM, Seumois G, Yates JR, Paulson JC, Bosinger SE, Wilson IA, Schief WR, Crotty S. The human naive b cell repertoire contains distinct subclasses for a germline-targeting hiv-1 vaccine immunogen. Sci Transl Med. 2018;10(448):10.1126/scitranslmed.aat0381

3. Xiao X, Chen W, Feng Y, Zhu Z, Prabakaran P, Wang Y, Zhang MY, Longo NS, Dimitrov DS. Germline-like predecessors of broadly neutralizing antibodies lack measurable binding to hiv-1 envelope glycoproteins: Implications for evasion of immune responses and design of vaccine immunogens. Biochem Biophys Res Commun. 2009;390(3):404–409. 10.1016/j.bbrc.2009.09.029

4. Xiao X, Chen W, Feng Y, Dimitrov DS. Maturation pathways of cross-reactive hiv-1 neutralizing antibodies. Viruses. 2009;1(3):802–817. 10.3390/v1030802

5. Wu X, Zhou T, Zhu J, Zhang B, Georgiev I, Wang C, Chen X, Longo NS, Louder M, McKee K, O’Dell S, Perfetto S, Schmidt SD, Shi W, Wu L, Yang Y, Yang ZY, Yang Z, Zhang Z, Bonsignori M, Crump JA, Kapiga SH, Sam NE, Haynes BF, Simek M, Burton DR, Koff WC, Doria-Rose NA, Connors M, Program NCS, Mullikin JC, Nabel GJ, Roederer M, Shapiro L, Kwong PD, Mascola JR. Focused evolution of hiv-1 neutralizing antibodies revealed by structures and deep sequencing. Science. 2011;333(6049):1593–1602. 10.1126/science.1207532

6. Klein F, Diskin R, Scheid JF, Gaebler C, Mouquet H, Georgiev IS, Pancera M, Zhou T, Incesu RB, Fu BZ, Gnanapragasam PN, Oliveira TY, Seaman MS, Kwong PD, Bjorkman PJ, Nussenzweig MC. Somatic mutations of the immunoglobulin framework are generally required for broad and potent hiv-1 neutralization. Cell. 2013;153(1):126–138. 10.1016/j.cell.2013.03.018

7. Liao HX, Lynch R, Zhou T, Gao F, Alam SM, Boyd SD, Fire AZ, Roskin KM, Schramm CA, Zhang Z, Zhu J, Shapiro L, Program NCS, Mullikin JC, Gnanakaran S, Hraber P, Wiehe K, Kelsoe G, Yang G, Xia SM, Montefiori DC, Parks R, Lloyd KE, Scearce RM, Soderberg KA, Cohen M, Kamanga G, Louder MK, Tran LM, Chen Y, Cai F, Chen S, Moquin S, Du X, Joyce MG, Srivatsan S, Zhang B, Zheng A, Shaw GM, Hahn BH, Kepler TB, Korber BT, Kwong PD, Mascola JR, Haynes BF. Co-evolution of a broadly neutralizing hiv-1 antibody and founder virus. Nature. 2013;496(7446):469–476. 10.1038/nature12053

8. Wiehe K, Bradley T, Meyerhoff RR, Hart C, Williams WB, Easterhoff D, Faison WJ, Kepler TB, Saunders KO, Alam SM, Bonsignori M, Haynes BF. Functional relevance of improbable antibody mutations for hiv broadly neutralizing antibody development. Cell Host Microbe. 2018;23(6):759–765 e756. 10.1016/j.chom.2018.04.018

9. Haynes BF, Fleming J, St Clair EW, Katinger H, Stiegler G, Kunert R, Robinson J, Scearce RM, Plonk K, Staats HF, Ortel TL, Liao HX, Alam SM. Cardiolipin polyspecific autoreactivity in two broadly neutralizing hiv-1 antibodies. Science. 2005;308(5730):1906–1908. 10.1126/science.1111781

10. Haynes BF, Wiehe K, Borrow P, Saunders KO, Korber B, Wagh K, McMichael AJ, Kelsoe G, Hahn BH, Alt F, Shaw GM. Strategies for hiv-1 vaccines that induce broadly neutralizing antibodies. Nat Rev Immunol. 2023;23(3):142–158. 10.1038/s41577-022-00753-w

11. Mu Z, Haynes BF, Cain DW. Strategies for eliciting multiple lineages of broadly neutralizing antibodies to hiv by vaccination. Curr Opin Virol. 2021;51(172-178. 10.1016/j.coviro.2021.09.015

12. Haynes BF, Kelsoe G, Harrison SC, Kepler TB. B-cell-lineage immunogen design in vaccine development with hiv-1 as a case study. Nat Biotechnol. 2012;30(5):423–433. 10.1038/nbt.2197

13. Wiehe K, Saunders KO, Stalls V, Cain DW, Venkatayogi S, Martin Beem JS, Berry M, Evangelous T, Henderson R, Hora B, Xia SM, Jiang C, Newman A, Bowman C, Lu X, Bryan ME, Bal J, Sanzone A, Chen H, Eaton A, Tomai MA, Fox CB, Tam YK, Barbosa C, Bonsignori M, Muramatsu H, Alam SM, Montefiori DC, Williams WB, Pardi N, Tian M, Weissman D, Alt FW, Acharya P, Haynes BF. Mutation-guided vaccine design: A process for developing boosting immunogens for hiv broadly neutralizing antibody induction. Cell Host Microbe. 2024;32(5):693–709 e697. 10.1016/j.chom.2024.04.006

14. Steichen JM, Lin YC, Havenar-Daughton C, Pecetta S, Ozorowski G, Willis JR, Toy L, Sok D, Liguori A, Kratochvil S, Torres JL, Kalyuzhniy O, Melzi E, Kulp DW, Raemisch S, Hu X, Bernard SM, Georgeson E, Phelps N, Adachi Y, Kubitz M, Landais E, Umotoy J, Robinson A, Briney B, Wilson IA, Burton DR, Ward AB, Crotty S, Batista FD, Schief WR. A generalized hiv vaccine design strategy for priming of broadly neutralizing antibody responses. Science. 2019;366(6470):10.1126/science.aax4380

15. Saunders KO, Wiehe K, Tian M, Acharya P, Bradley T, Alam SM, Go EP, Scearce R, Sutherland L, Henderson R, Hsu AL, Borgnia MJ, Chen H, Lu X, Wu NR, Watts B, Jiang C, Easterhoff D, Cheng HL, McGovern K, Waddicor P, Chapdelaine-Williams A, Eaton A, Zhang J, Rountree W, Verkoczy L, Tomai M, Lewis MG, Desaire HR, Edwards RJ, Cain DW, Bonsignori M, Montefiori D, Alt FW, Haynes BF. Targeted selection of hiv-specific antibody mutations by engineering b cell maturation. Science. 2019;366(6470):10.1126/science.aay7199

16. Jardine JG, Ota T, Sok D, Pauthner M, Kulp DW, Kalyuzhniy O, Skog PD, Thinnes TC, Bhullar D, Briney B, Menis S, Jones M, Kubitz M, Spencer S, Adachi Y, Burton DR, Schief WR, Nemazee D. Hiv-1 vaccines. Priming a broadly neutralizing antibody response to hiv-1 using a germline-targeting immunogen. Science. 2015;349(6244):156–161. 10.1126/science.aac5894

17. Huang D, Abbott RK, Havenar-Daughton C, Skog PD, Al-Kolla R, Groschel B, Blane TR, Menis S, Tran JT, Thinnes TC, Volpi SA, Liguori A, Schiffner T, Villegas SM, Kalyuzhniy O, Pintea M, Voss JE, Phelps N, Tingle R, Rodriguez AR, Martin G, Kupryianov S, deCamp A, Schief WR, Nemazee D, Crotty S. B cells expressing authentic naive human vrc01-class bcrs can be recruited to germinal centers and affinity mature in multiple independent mouse models. Proc Natl Acad Sci U S A. 2020;117(37):22920–22931. 10.1073/pnas.2004489117

18. Caniels TG, Medina-Ramirez M, Zhang J, Sarkar A, Kumar S, LaBranche A, Derking R, Allen JD, Snitselaar JL, Capella-Pujol J, Sanchez IDM, Yasmeen A, Diaz M, Aldon Y, Bijl TPL, Venkatayogi S, Martin Beem JS, Newman A, Jiang C, Lee WH, Pater M, Burger JA, van Breemen MJ, de Taeye SW, Rantalainen K, LaBranche C, Saunders KO, Montefiori D, Ozorowski G, Ward AB, Crispin M, Moore JP, Klasse PJ, Haynes BF, Wilson IA, Wiehe K, Verkoczy L, Sanders RW. Germline-targeting hiv-1 env vaccination induces vrc01-class antibodies with rare insertions. Cell Rep Med. 2023;4(4):101003. 10.1016/j.xcrm.2023.101003

19. Leggat DJ, Cohen KW, Willis JR, Fulp WJ, deCamp AC, Kalyuzhniy O, Cottrell CA, Menis S, Finak G, Ballweber-Fleming L, Srikanth A, Plyler JR, Schiffner T, Liguori A, Rahaman F, Lombardo A, Philiponis V, Whaley RE, Seese A, Brand J, Ruppel AM, Hoyland W, Yates NL, Williams LD, Greene K, Gao H, Mahoney CR, Corcoran MM, Cagigi A, Taylor A, Brown DM, Ambrozak DR, Sincomb T, Hu X, Tingle R, Georgeson E, Eskandarzadeh S, Alavi N, Lu D, Mullen TM, Kubitz M, Groschel B, Maenza J, Kolokythas O, Khati N, Bethony J, Crotty S, Roederer M, Karlsson Hedestam GB, Tomaras GD, Montefiori D, Diemert D, Koup RA, Laufer DS, McElrath MJ, McDermott AB, Schief WR. Vaccination induces hiv broadly neutralizing antibody precursors in humans. Science. 2022;378(6623):eadd6502. 10.1126/science.add6502

20. Williams WB, Alam SM, Ofek G, Erdmann N, Montefiori DC, Seaman MS, Wagh K, Korber B, Edwards RJ, Mansouri K, Eaton A, Cain DW, Martin M, Hwang J, Arus-Altuz A, Lu X, Cai F, Jamieson N, Parks R, Barr M, Foulger A, Anasti K, Patel P, Sammour S, Parsons RJ, Huang X, Lindenberger J, Fetics S, Janowska K, Niyongabo A, Janus BM, Astavans A, Fox CB, Mohanty I, Evangelous T, Chen Y, Berry M, Kirshner H, Van Itallie E, Saunders KO, Wiehe K, Cohen KW, McElrath MJ, Corey L, Acharya P, Walsh SR, Baden LR, Haynes BF. Vaccine induction of heterologous hiv-1-neutralizing antibody b cell lineages in humans. Cell. 2024;187(12):2919–2934 e2920. 10.1016/j.cell.2024.04.033

21. Bachmann MF, Odermatt B, Hengartner H, Zinkernagel RM. Induction of long-lived germinal centers associated with persisting antigen after viral infection. J Exp Med. 1996;183(5):2259–2269. 10.1084/jem.183.5.2259

22. Yewdell WT, Smolkin RM, Belcheva KT, Mendoza A, Michaels AJ, Cols M, Angeletti D, Yewdell JW, Chaudhuri J. Temporal dynamics of persistent germinal centers and memory b cell differentiation following respiratory virus infection. Cell Rep. 2021;37(6):109961. 10.1016/j.celrep.2021.109961

23. de Carvalho RVH, Ersching J, Barbulescu A, Hobbs A, Castro TBR, Mesin L, Jacobsen JT, Phillips BK, Hoffmann HH, Parsa R, Canesso MCC, Nowosad CR, Feng A, Leist SR, Baric RS, Yang E, Utz PJ, Victora GD. Clonal replacement sustains long-lived germinal centers primed by respiratory viruses. Cell. 2023;186(1):131–146 e113. 10.1016/j.cell.2022.11.031

24. Hagglof T, Cipolla M, Loewe M, Chen ST, Mesin L, Hartweger H, ElTanbouly MA, Cho A, Gazumyan A, Ramos V, Stamatatos L, Oliveira TY, Nussenzweig MC, Viant C. Continuous germinal center invasion contributes to the diversity of the immune response. Cell. 2023;186(1):147–161 e115. 10.1016/j.cell.2022.11.032

25. Mesin L, Schiepers A, Ersching J, Barbulescu A, Cavazzoni CB, Angelini A, Okada T, Kurosaki T, Victora GD. Restricted clonality and limited germinal center reentry characterize memory b cell reactivation by boosting. Cell. 2020;180(1):92–106 e111. 10.1016/j.cell.2019.11.032

26. Zabel F, Mohanan D, Bessa J, Link A, Fettelschoss A, Saudan P, Kundig TM, Bachmann MF. Viral particles drive rapid differentiation of memory b cells into secondary plasma cells producing increased levels of antibodies. J Immunol. 2014;192(12):5499–5508. 10.4049/jimmunol.1400065

27. Pape KA, Taylor JJ, Maul RW, Gearhart PJ, Jenkins MK. Different b cell populations mediate early and late memory during an endogenous immune response. Science. 2011;331(6021):1203–1207. 10.1126/science.1201730

28. Dogan I, Bertocci B, Vilmont V, Delbos F, Megret J, Storck S, Reynaud CA, Weill JC. Multiple layers of b cell memory with different effector functions. Nat Immunol. 2009;10(12):1292–1299. 10.1038/ni.1814

29. Phan TG, Paus D, Chan TD, Turner ML, Nutt SL, Basten A, Brink R. High affinity germinal center b cells are actively selected into the plasma cell compartment. J Exp Med. 2006;203(11):2419–2424. 10.1084/jem.20061254

30. Paus D, Phan TG, Chan TD, Gardam S, Basten A, Brink R. Antigen recognition strength regulates the choice between extrafollicular plasma cell and germinal center b cell differentiation. J Exp Med. 2006;203(4):1081–1091. 10.1084/jem.20060087

31. Allie SR, Bradley JE, Mudunuru U, Schultz MD, Graf BA, Lund FE, Randall TD. The establishment of resident memory b cells in the lung requires local antigen encounter. Nat Immunol. 2019;20(1):97–108. 10.1038/s41590-018-0260-6

32. Kuraoka M, Yeh CH, Bajic G, Kotaki R, Song S, Windsor I, Harrison SC, Kelsoe G. Recall of b cell memory depends on relative locations of prime and boost immunization. Sci Immunol. 2022;7(71):eabn5311. 10.1126/sciimmunol.abn5311

33. Dhenni R, Hoppe AC, Reynaldi A, Kyaw W, Handoko NT, Grootveld AK, Keith YH, Bhattacharyya ND, Ahel HI, Telfser AJ, McCorkindale AN, Yazar S, Bui CHT, Smith JT, Khoo WH, Boyd M, Obeid S, Milner B, Starr M, Brilot F, Milogiannakis V, Akerman A, Aggarwal A, Davenport MP, Deenick EK, Chaffer CL, Croucher PI, Brink R, Goldstein LD, Cromer D, Turville SG, Kelleher AD, Venturi V, Munier CML, Phan TG. Macrophages direct location-dependent recall of b cell memory to vaccination. Cell. 2025;188(13):3477–3496 e3422. 10.1016/j.cell.2025.04.005

34. Zuccarino-Catania GV, Sadanand S, Weisel FJ, Tomayko MM, Meng H, Kleinstein SH, Good-Jacobson KL, Shlomchik MJ. Cd80 and pd-l2 define functionally distinct memory b cell subsets that are independent of antibody isotype. Nat Immunol. 2014;15(7):631–637. 10.1038/ni.2914

35. Adachi Y, Onodera T, Yamada Y, Daio R, Tsuiji M, Inoue T, Kobayashi K, Kurosaki T, Ato M, Takahashi Y. Distinct germinal center selection at local sites shapes memory b cell response to viral escape. J Exp Med. 2015;212(10):1709–1723. 10.1084/jem.20142284

36. Shinnakasu R, Inoue T, Kometani K, Moriyama S, Adachi Y, Nakayama M, Takahashi Y, Fukuyama H, Okada T, Kurosaki T. Regulated selection of germinal-center cells into the memory b cell compartment. Nat Immunol. 2016;17(7):861–869. 10.1038/ni.3460

37. Kuraoka M, Schmidt AG, Nojima T, Feng F, Watanabe A, Kitamura D, Harrison SC, Kepler TB, Kelsoe G. Complex antigens drive permissive clonal selection in germinal centers. Immunity. 2016;44(3):542–552. 10.1016/j.immuni.2016.02.010

38. Bonsignori M, Kreider EF, Fera D, Meyerhoff RR, Bradley T, Wiehe K, Alam SM, Aussedat B, Walkowicz WE, Hwang KK, Saunders KO, Zhang R, Gladden MA, Monroe A, Kumar A, Xia SM, Cooper M, Louder MK, McKee K, Bailer RT, Pier BW, Jette CA, Kelsoe G, Williams WB, Morris L, Kappes J, Wagh K, Kamanga G, Cohen MS, Hraber PT, Montefiori DC, Trama A, Liao HX, Kepler TB, Moody MA, Gao F, Danishefsky SJ, Mascola JR, Shaw GM, Hahn BH, Harrison SC, Korber BT, Haynes BF. Staged induction of hiv-1 glycan-dependent broadly neutralizing antibodies. Sci Transl Med. 2017;9(381):10.1126/scitranslmed.aai7514

39. Montefiori DC. Measuring hiv neutralization in a luciferase reporter gene assay. Methods Mol Biol. 2009;485(395-405. 10.1007/978-1-59745-170-3_26

40. Li M, Gao F, Mascola JR, Stamatatos L, Polonis VR, Koutsoukos M, Voss G, Goepfert P, Gilbert P, Greene KM, Bilska M, Kothe DL, Salazar-Gonzalez JF, Wei X, Decker JM, Hahn BH, Montefiori DC. Human immunodeficiency virus type 1 env clones from acute and early subtype b infections for standardized assessments of vaccine-elicited neutralizing antibodies. J Virol. 2005;79(16):10108–10125. 10.1128/JVI.79.16.10108-10125.2005

41. Stuart T, Butler A, Hoffman P, Hafemeister C, Papalexi E, Mauck WM, 3rd, Hao Y, Stoeckius M, Smibert P, Satija R. Comprehensive integration of single-cell data. Cell. 2019;177(7):1888–1902 e1821. 10.1016/j.cell.2019.05.031

42. Yeh CH, Finney J, Okada T, Kurosaki T, Kelsoe G. Primary germinal center-resident t follicular helper cells are a physiologically distinct subset of cxcr5(hi)pd-1(hi) t follicular helper cells. Immunity. 2022;55(2):272–289 e277. 10.1016/j.immuni.2021.12.015

43. Schiepers A, van ’t Wout MFL, Greaney AJ, Zang T, Muramatsu H, Lin PJC, Tam YK, Mesin L, Starr TN, Bieniasz PD, Pardi N, Bloom JD, Victora GD. Molecular fate-mapping of serum antibody responses to repeat immunization. Nature. 2023;615(7952):482–489. 10.1038/s41586-023-05715-3

44. Chernova I, Jones DD, Wilmore JR, Bortnick A, Yucel M, Hershberg U, Allman D. Lasting antibody responses are mediated by a combination of newly formed and established bone marrow plasma cells drawn from clonally distinct precursors. J Immunol. 2014;193(10):4971–4979. 10.4049/jimmunol.1401264

45. Klasse PJ, Ozorowski G, Sanders RW, Moore JP. Env exceptionalism: Why are hiv-1 env glycoproteins atypical immunogens? Cell Host Microbe. 2020;27(4):507–518. 10.1016/j.chom.2020.03.018

46. Cyster JG, Wilson PC. Antibody modulation of b cell responses-incorporating positive and negative feedback. Immunity. 2024;57(7):1466–1481. 10.1016/j.immuni.2024.06.009

47. Laidlaw BJ, Duan L, Xu Y, Vazquez SE, Cyster JG. The transcription factor hhex cooperates with the corepressor tle3 to promote memory b cell development. Nat Immunol. 2020;21(9):1082–1093. 10.1038/s41590-020-0713-6

48. Pikor NB, Morbe U, Lutge M, Gil-Cruz C, Perez-Shibayama C, Novkovic M, Cheng HW, Nombela-Arrieta C, Nagasawa T, Linterman MA, Onder L, Ludewig B. Remodeling of light and dark zone follicular dendritic cells governs germinal center responses. Nat Immunol. 2020;21(6):649–659. 10.1038/s41590-020-0672-y

49. Victora GD, Dominguez-Sola D, Holmes AB, Deroubaix S, Dalla-Favera R, Nussenzweig MC. Identification of human germinal center light and dark zone cells and their relationship to human b-cell lymphomas. Blood. 2012;120(11):2240–2248. 10.1182/blood-2012-03-415380

50. Inoue T, Kurosaki T. Memory b cells. Nat Rev Immunol. 2024;24(1):5-17. 10.1038/s41577-023-00897-3

51. Schiepers A, Van’t Wout MFL, Hobbs A, Mesin L, Victora GD. Opposing effects of pre-existing antibody and memory t cell help on the dynamics of recall germinal centers. Immunity. 2024;57(7):1618–1628 e1614. 10.1016/j.immuni.2024.05.009

52. Inoue T, Moran I, Shinnakasu R, Phan TG, Kurosaki T. Generation of memory b cells and their reactivation. Immunol Rev. 2018;283(1):138–149. 10.1111/imr.12640

53. Turner JS, Zhou JQ, Han J, Schmitz AJ, Rizk AA, Alsoussi WB, Lei T, Amor M, McIntire KM, Meade P, Strohmeier S, Brent RI, Richey ST, Haile A, Yang YR, Klebert MK, Suessen T, Teefey S, Presti RM, Krammer F, Kleinstein SH, Ward AB, Ellebedy AH. Human germinal centres engage memory and naive b cells after influenza vaccination. Nature. 2020;586(7827):127–132. 10.1038/s41586-020-2711-0

54. Kim W, Zhou JQ, Horvath SC, Schmitz AJ, Sturtz AJ, Lei T, Liu Z, Kalaidina E, Thapa M, Alsoussi WB, Haile A, Klebert MK, Suessen T, Parra-Rodriguez L, Mudd PA, Whelan SPJ, Middleton WD, Teefey SA, Pusic I, O’Halloran JA, Presti RM, Turner JS, Ellebedy AH. Germinal centre-driven maturation of b cell response to mrna vaccination. Nature. 2022;604(7904):141–145. 10.1038/s41586-022-04527-1

55. Lee JH, Sutton HJ, Cottrell CA, Phung I, Ozorowski G, Sewall LM, Nedellec R, Nakao C, Silva M, Richey ST, Torres JL, Lee WH, Georgeson E, Kubitz M, Hodges S, Mullen TM, Adachi Y, Cirelli KM, Kaur A, Allers C, Fahlberg M, Grasperge BF, Dufour JP, Schiro F, Aye PP, Kalyuzhniy O, Liguori A, Carnathan DG, Silvestri G, Shen X, Montefiori DC, Veazey RS, Ward AB, Hangartner L, Burton DR, Irvine DJ, Schief WR, Crotty S. Long-primed germinal centres with enduring affinity maturation and clonal migration. Nature. 2022;609(7929):998–1004. 10.1038/s41586-022-05216-9

56. Cirelli KM, Carnathan DG, Nogal B, Martin JT, Rodriguez OL, Upadhyay AA, Enemuo CA, Gebru EH, Choe Y, Viviano F, Nakao C, Pauthner MG, Reiss S, Cottrell CA, Smith ML, Bastidas R, Gibson W, Wolabaugh AN, Melo MB, Cossette B, Kumar V, Patel NB, Tokatlian T, Menis S, Kulp DW, Burton DR, Murrell B, Schief WR, Bosinger SE, Ward AB, Watson CT, Silvestri G, Irvine DJ, Crotty S. Slow delivery immunization enhances hiv neutralizing antibody and germinal center responses via modulation of immunodominance. Cell. 2019;177(5):1153–1171 e1128. 10.1016/j.cell.2019.04.012

57. Uhr JW, Moller G. Regulatory effect of antibody on the immune response. Adv Immunol. 1968;8(81-127. 10.1016/s0065-2776(08)60465-4

58. Tas JMJ, Koo JH, Lin YC, Xie Z, Steichen JM, Jackson AM, Hauser BM, Wang X, Cottrell CA, Torres JL, Warner JE, Kirsch KH, Weldon SR, Groschel B, Nogal B, Ozorowski G, Bangaru S, Phelps N, Adachi Y, Eskandarzadeh S, Kubitz M, Burton DR, Lingwood D, Schmidt AG, Nair U, Ward AB, Schief WR, Batista FD. Antibodies from primary humoral responses modulate the recruitment of naive b cells during secondary responses. Immunity. 2022;55(10):1856–1871 e1856. 10.1016/j.immuni.2022.07.020

59. Zhang Y, Meyer-Hermann M, George LA, Figge MT, Khan M, Goodall M, Young SP, Reynolds A, Falciani F, Waisman A, Notley CA, Ehrenstein MR, Kosco-Vilbois M, Toellner KM. Germinal center b cells govern their own fate via antibody feedback. J Exp Med. 2013;210(3):457–464. 10.1084/jem.20120150

60. Inoue T, Shinnakasu R, Kawai C, Yamamoto H, Sakakibara S, Ono C, Itoh Y, Terooatea T, Yamashita K, Okamoto T, Hashii N, Ishii-Watabe A, Butler NS, Matsuura Y, Matsumoto H, Otsuka S, Hiraoka K, Teshima T, Murakami M, Kurosaki T. Antibody feedback contributes to facilitating the development of omicron-reactive memory b cells in sars-cov-2 mrna vaccinees. J Exp Med. 2023;220(2):10.1084/jem.20221786

61. Roozendaal R, Mempel TR, Pitcher LA, Gonzalez SF, Verschoor A, Mebius RE, von Andrian UH, Carroll MC. Conduits mediate transport of low-molecular-weight antigen to lymph node follicles. Immunity. 2009;30(2):264–276. 10.1016/j.immuni.2008.12.014

62. Phan TG, Grigorova I, Okada T, Cyster JG. Subcapsular encounter and complement-dependent transport of immune complexes by lymph node b cells. Nat Immunol. 2007;8(9):992–1000. 10.1038/ni1494

